# The role of DDK and Treslin-MTBP in coordinating replication licensing and pre-Initiation Complex formation

**DOI:** 10.1101/2021.03.22.436442

**Authors:** Ilaria Volpi, Peter J. Gillespie, Gaganmeet Singh Chadha, J. Julian Blow

## Abstract

Treslin/Ticrr is required for the initiation of DNA replication and binds to MTBP (Mdm2 Binding Protein). Here we show that in *Xenopus* egg extract, MTBP forms an elongated tetramer with Treslin containing two molecules of each protein. Immunodepletion and add-back experiments show that Treslin-MTBP is rate-limiting for replication initiation. It is recruited onto chromatin before S phase starts and recruitment continues during S phase. We show that DDK activity both increases and strengthens the interaction of Treslin-MTBP with licensed chromatin. We also show that DDK activity cooperates with CDK activity to drive the interaction of Treslin-MTBP with TopBP1 which is a regulated crucial step in pre-Initiation Complex formation. These results suggest how DDK works together with CDKs to regulate Treslin-MTBP and plays a crucial in selecting which origins will undergo initiation.

## Introduction

The initiation of eukaryotic DNA replication occurs at replication origins that have been licensed by being loaded with double hexamers of MCM2-7 proteins. The combined action of two kinases, DDK and S-CDK, then triggers the recruitment of Cdc45 and the GINS complex onto these MCM2-7 double hexamers to form the replicative CMG (Cdc45-MCM-GINS) helicase. *In vitro* experiments in *Xenopus* egg extract have shown that DDK activity is required before S-CDK activity (Jares and Blow, 2000; Walter, 2000). DDK-mediated hyperphosphorylation of Mcm4 but not Mcm2 correlates with replication initiation in both *Xenopus* egg extract and human cells (Poh et al., 2014; Alver et al., 2017). In *S. cerevisiae*, DDK phosphorylation of MCM2-7 promotes the recruitment of Sld3 and Cdc45 to origins, and then S-CDK phosphorylation of Sld3 and Sld2 allows the recruitment of Dbp11, Sld2, Sld3, GINS, and Pol ε (Muramatsu et al., 2010; Li and Araki, 2013; Yeeles et al., 2015; Deegan et al., 2016). DDK-dependent MCM phosphorylation is reversed by protein phosphatase 1 targeted to chromatin by Rif1 in *Xenopus* egg extract and in both *S. cerevisiae* and human cells (Alver *et al*., 2017; Hiraga et al., 2014; Hiraga et al., 2017; Poh *et al*., 2014).

Sld7 was identified in *S. cerevisiae* as a gene that is synthetically lethal with Dpb11 (Kamimura et al., 1998; Tanaka et al., 2011a). Despite Sld7 not being absolutely required for DNA replication, it physically interacts with Sld3 and is required for the proper function of Sld3 at initiation (Deegan *et al*., 2016; Tanaka *et al*., 2011a; Tanaka et al., 2011b). Sld7 mutants have reduced total levels of Sld3 and delayed dissociation of Sld3 from GINS at replication initiation (Tanaka *et al*., 2011a). The crystal structures obtained from Sld3 and Sld7 domains, suggest a model of interaction between these proteins in which two molecules of Sld7 interact with each other in an antiparallel manner through their C-terminal domains (Itou et al., 2015). The N-terminal domain of Sld7 interacts with the N-terminal domain of Sld3 to form a tetramer in which the two molecules of Sld3 are connected by the two molecules of Sld7 (Itou *et al*., 2015). This conformation could allow the correct positioning of Sld3 onto the double hexamer of MCM2-7 in order to promote bidirectional firing of the origins.

MTBP was identified in a yeast two-hybrid screen as an interacting partner of MDM2, the E3 ubiquitin ligase that targets p53 for destruction (Boyd et al., 2000). In human cells, knockdown of MTBP blocks cell growth and induces a G1 arrest (Boyd *et al*., 2000; Odvody et al., 2010). It is frequently found overexpressed in cancer (Grieb et al., 2014b; Grieb et al., 2014a). In human cells MTBP has been identified as an interacting partner of Treslin, the metazoan homologue of Sld3, throughout the cell cycle (Boos et al., 2013). An MTBP interacting domain has been identified on Treslin, upstream of the Cdc45- interacting domain (Boos *et al*., 2013). Treslin lacking the MTBP-interacting domain is unable to rescue the DNA replication phenotype of Treslin-depleted cells, indicating that the Treslin-MTBP interaction is essential for replication (Boos *et al*., 2013). MTBP siRNA causes the impairment of CMG assembly and the lengthening of S phase, suggesting a direct role in the initiation of DNA replication (Boos *et al*., 2013).

Treslin interacts with TopBP1, the metazoan homologue of Dpb11. CDK-dependent phosphorylation of the well-conserved S1001 of Treslin mediates its interaction with the phospho-binding BRCT domains of TopBP1 (Kumagai et al., 2011; Boos *et al*., 2013). Mutation of this residue, but not of the adjacent well-conserved CDK consensus-site T969, significantly reduced the interaction of Treslin with TopBP1 and cells harbouring a non-phosphorylatable mutation of S1001 were deficient in DNA replication (Kumagai *et al*., 2011). The formation of a tripartite Treslin-MTBP-TopBP1 complex in human cells is curtailed when Treslin’s conserved phosphosites are mutated (Boos *et al*., 2013). CDK phosphorylation of Treslin is therefore required both for the interaction of Treslin-MTBP with TopBP1 and to support DNA replication in human cells. Consistent with this, cells with a phosphomimetic Treslin mutant display faster replication kinetics and a shorter S phase (Sansam et al., 2015). In addition to Treslin-MTBP, RecQ4, the higher eukaryotic homologue of yeast Sld2, is a potential CDK substrate. Following phosphorylation of RecQ4 it associates with licensed chromatin, together with Treslin-MTBP, to form the pre-Initiation complex (preIC) (Im et al., 2015).

Here we further characterise the role of MTBP in DNA replication using the *Xenopus* egg extract cell-free system. Our work complements and extends a recent report describing *Xenopus* MTBP (Kumagai and Dunphy, 2017). We show that MTBP and Treslin form a tetrameric complex that is required for the initiation of DNA replication, similar to Sld3/Sld7 in yeast. However, in this system, the regulation of MTBP binding onto chromatin appears substantially different from Sld7 in yeast. We find that DDK activity is required to both to increase and strengthen the interaction of Treslin-MTBP with chromatin and to mediate its association with TopBP1. Our data suggest a key role for DDK activity in co-ordinating the interaction of Treslin-MTBP with licensed replication origins and TopBP1, thereby determining which origins are selected to undergo replication initiation.

## Results

### Characterisation of MTBP and Treslin antibodies

In human cells MTBP has been identified as an interacting partner of Treslin involved in regulation of the initiation of DNA replication(Boos *et al*., 2013). To identify how MTBP is involved in DNA replication we studied its role and regulation in *Xenopus* egg extracts which support complete genome duplication *in vitro* (Gillespie et al., 2016). We raised antibodies against the N’ and C’ termini of *Xenopus* MTBP, which we denote MTBP-1 and MTBP-2, respectively. We also used a commercial antibody raised against a conserved central region of the human protein, which we denote by MTBP-S. The antibodies were used to immunoblot whole *Xenopus* egg extract, replicating chromatin, and immunoprecipitates (Fig. S1a). Whilst all the antibodies detected multiple bands in whole extract, they each recognised the same single band on replicating chromatin of the expected molecular weight for MTBP. Importantly, they each also specifically immunoprecipitated the same band. We also raised two antibodies against *Xenopus* Treslin (Treslin-1 and Treslin-2), which showed a similar specificity (Fig. S1b) and an antibody against *Xenopus* TopBP1 which recognised only a single band of the expected Mw in both whole egg extract and on isolated chromatin Fig. S1c). To confirm the identity of the major band identified by the MTBP and Treslin antibodies, we performed a mass spectrometry of the immunoprecipitates, which confirmed that MTBP and Treslin were major components of these samples (Fig. S2).

### A Treslin-MTBP complex in Xenopus egg extract

To identify a possible Treslin-MTBP complex we immunoprecipitated Treslin and MTBP from aliquots of *Xenopus* egg extract in either metaphase or interphase (Fig. 1a). Immunoblotting showed that MTBP antibodies co-precipitated Treslin and Treslin antibodies co-precipitated MTBP. These results indicate that in *Xenopus* egg extract, MTBP and Treslin form a complex throughout the cell cycle. This is consistent with previous reports in *Xenopus* egg extracts (Kumagai and Dunphy, 2017) and human cells (Boos *et al*., 2013) and resembles the interaction between Sld3 and Sld7 in yeast (Tanaka *et al*., 2011a; Tanaka *et al*., 2011b).

**Figure 1.**
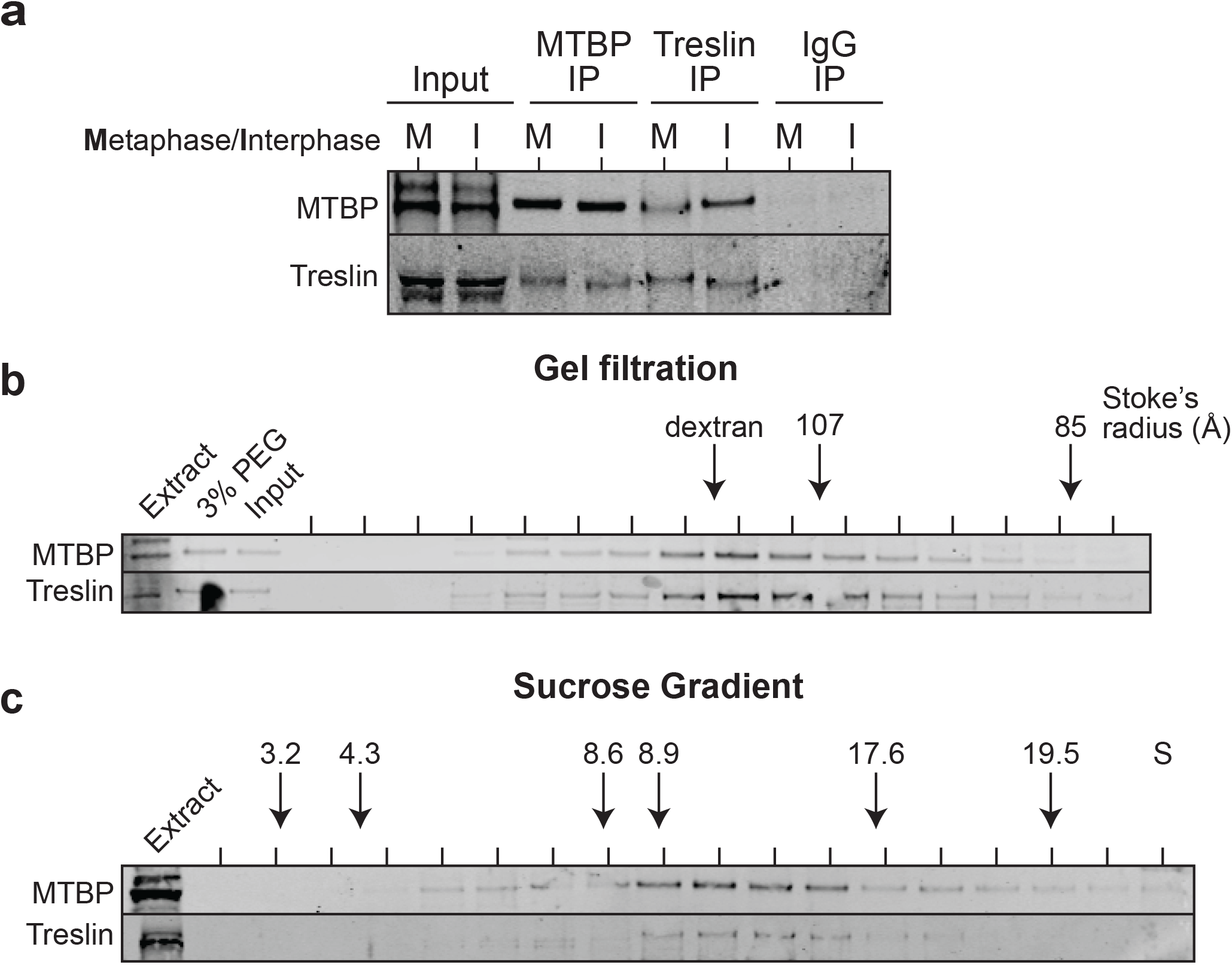
Treslin-MTBP complex in *Xenopus* egg extract. **a.** Immunoprecipitation of MTBP and Treslin from metaphase (M) or interphase (I) extract using affinity purified MTBP-1 and Treslin-1 antibodies, immunoblotted for MTBP and Treslin. **b.** Gel filtration of 3% PEG-precipitated proteins from *Xenopus* egg extract immunoblotted for MTBP and Treslin. **c.** 5-40% sucrose gradient centrifugation of 3%-PEG precipitated proteins, immunoblotted for MTBP and Treslin.

We next set out to determine the molecular weight of the Treslin-MTBP complex. Elongated proteins appear larger than spherical proteins of the same molecular weight when analysed by size exclusion chromatography, but appear smaller when analysed by sucrose gradient centrifugation. However, by combining data from both separation techniques (Stoke’s radius and sedimentation coefficient) and using the Siegel and Monty equation (Siegel and Monty, 1966), the native molecular weight of a protein or protein complex can be determined in a manner that is unaffected by its shape. We precipitated proteins from a clarified supernatant of whole egg extract using polyethylene glycol (PEG) (see Fig. 2), selected the protein fraction containing MTBP and Treslin, then analysed it by size exclusion chromatography and sucrose gradient centrifugation. MTBP and Treslin showed similar elution profiles in size exclusion chromatography and sucrose gradient centrifugation, suggesting that in egg extract they form a single multimeric species. This complex has a Stoke’s radius of 116 Å (Fig. 1b) and a sedimentation coefficient of 12.8 S (Fig. 1c). Applying the Siegel and Monty equation gives a molecular weight for the Treslin-MTBP complex of 626 kDa. This is close to (± <1%) the expected molecular weight of a 632 kDa tetramer composed of two molecules of MTBP (96 kDa) and two molecules of Treslin (220 kDa). The Treslin-MTBP complex appears larger on gel filtration than thyroglobulin tetramer (1,338 kDa, 107 Å) and smaller on sucrose gradients than β-amylase (200 kDa, 54 Å, 8.9 S), suggesting that Treslin-MTBP is highly elongated.

**Figure 2.**
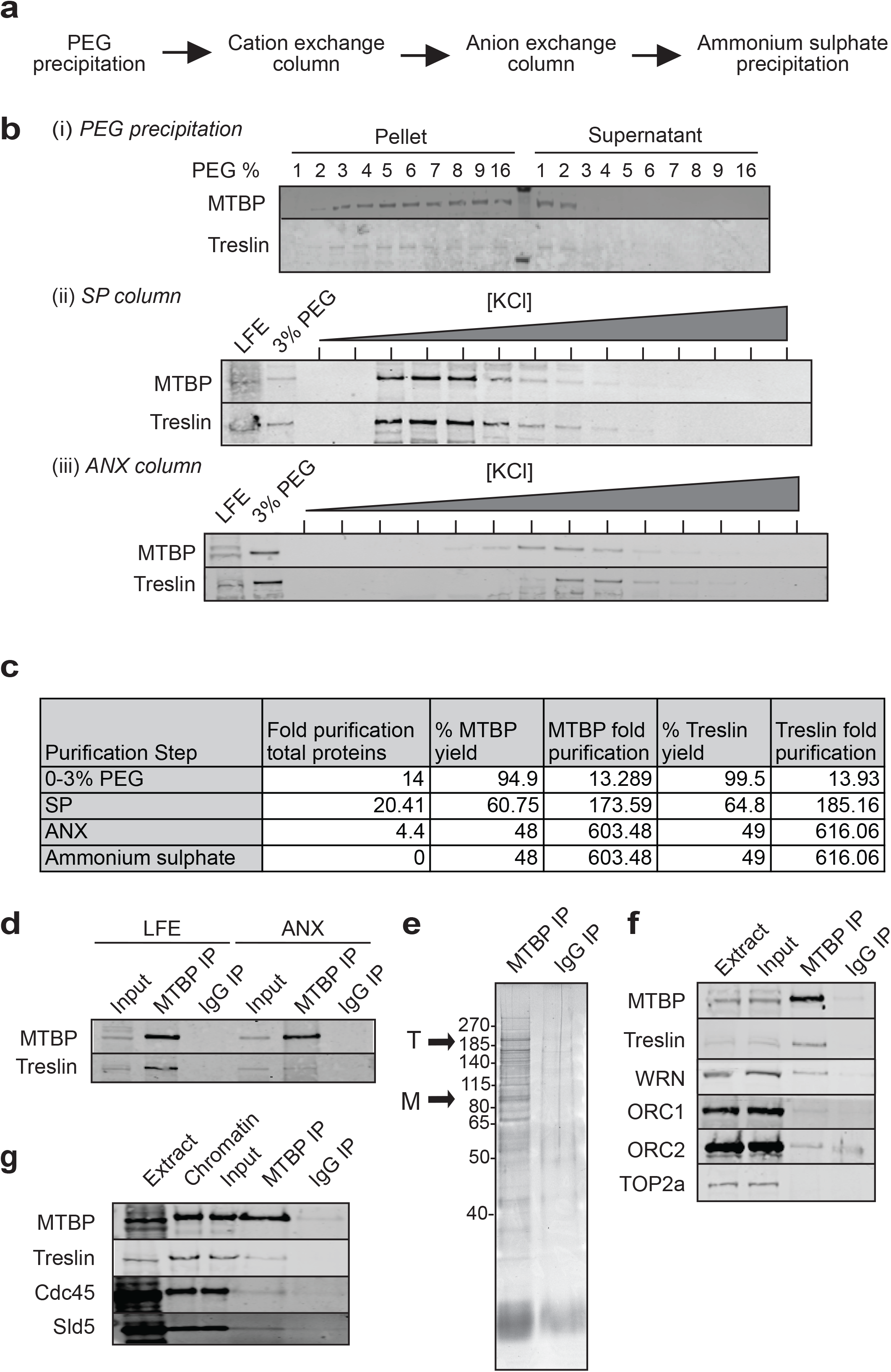
Treslin-MTBP complex purification. **a.** Flow chart of the purification process. **b**. Immunoblots of MTBP and Treslin present in (i) PEG, (ii) SP and (iii) ANX fractions. **c.** Table of purification achieved for MTBP and Treslin. **d, e.** Purified material was immunoprecipitated using MTBP-1 affinity purified antibody. Samples were immunoblotted for MTBP and Treslin (d) and run on a NuPage gel and stained with Sypro Ruby (e). **f.** MTBP immunoprecipitate from whole extract was run on NuPage gels and immunoblotted for the indicated proteins. **g.** Sperm nuclei were incubated in interphase extract for 30 min; chromatin was isolated, digested with DNAse and immunoprecipitated with MTBP-1 or non-immune antibodies. Immunoprecipitates were immunoblotted for the indicated proteins.

Quantitative immunoblotting that compares the signal of MTBP in extract with the signal from known quantities of the MTBP-2-antigen suggested that the concentration of MTBP in undiluted egg extract is ∼0.2 μg/ml or ∼2 nM (Fig. S3). The concentration of the proposed Treslin-MTBP tetramer is therefore ∼1 nM. *Xenopus* egg extract can efficiently replicate ∼30 ng DNA per microliter of extract (Blow and Laskey, 1986) with active replication origins spaced ∼10 kb apart (Blow et al., 2001). At 30 ng DNA/μl the concentration of active origins is therefore ∼4 nM, suggesting that Treslin-MTBP is involved in replication initiation at rate-limiting levels.

### Purification of Treslin-MTBP complex

To investigate the composition of the Treslin-MTBP complex we carried out a partial chromatographic purification (Fig. 2a). First, we precipitated proteins from a clarified supernatant of whole egg extract using PEG (Fig. 2B(i)), then used cation and anion exchange chromatography to further enrich MTBP and Treslin (Fig. 2B(ii-iii)). We carried out a final ammonium sulphate precipitation in order to concentrate the protein fractions. During the purification we achieved a 1200-fold reduction of total protein as estimated by UV absorbance at 280 nm with a yield of MTBP and Treslin of 48% and 49% respectively (Fig. 2c). Combining data for the yield with protein concentration measurements suggests that MTBP and Treslin were both enriched >600-fold (Fig. 2c).

Immunoprecipitation of the enriched material with MTBP-1 antibodies co-precipitated Treslin, though less efficiently than in whole extract, suggesting that the complex was only partially maintained throughout the chromatographic procedures (Fig. 2d); consistent with this a fraction of MTBP separate from Treslin is found on cation exchange chromatography (Fig. 2b(iii)). Total protein staining of the immunoprecipitates showed major bands at the expected position of MTBP and Treslin and as expected, the higher molecular weight Treslin band is the more intense (Fig. 2e). Mass spectrometry confirmed that MTBP and Treslin were major protein components of the immunoprecipitate (Fig. S4). We were unsuccessful in reducing the number of bands in the immunoprecipitates by increasing salt concentration during the wash steps, because this strongly reduced the co-precipitation of Treslin with MTBP antibodies. Taken together, our results strongly suggest that MTBP and Treslin form a 2:2 heterodimer, though the partial purification falls short of providing definitive evidence for this.

Mass spectrometry of the MTBP immunoprecipitate identified other proteins, including components of the Origin Recognition Complex (ORC), Werner’s helicase (WRN) and Topoisomerase IIα. To test whether any of these interactions were specific, we immunoblotted MTBP-1 immunoprecipitates with antibodies to these proteins. Orc1, Orc2 and WRN were detected in MTBP-1 immunoprecipitates (Fig. 2f). We therefore conclude that although MTBP and Treslin exist primarily as a hetero-tetramer in egg extract, they can also associate with other proteins to a limited extent.

The Treslin-MTBP hetero-tetramer is present in interphase egg extracts (Fig. 1). Following nuclear formation these extracts support one complete round of DNA replication. To determine whether the Treslin-MTBP complex is present on chromatin during S phase we performed a chromatin immunoprecipitation (ChIP) from replicating nuclei isolated from egg extract (Fig. 2g). We digested replicating chromatin with nuclease and performed MTBP immunoprecipitation. Treslin was also co-precipitated with MTBP, at levels comparable to the total amount loaded onto chromatin, suggesting that the two proteins remain associated with one another when bound to DNA. This also suggests that, consistent with the chromatography data, there is no significant excess of Treslin over MTBP in the extract. MTBP also co-precipitated Cdc45 and Sld5 on chromatin (Fig. 2g); these proteins are members of the CMG helicase, consistent with the idea that the Treslin-MTBP complex is involved in assembling the CMG complex.

### Regulation of MTBP recruitment onto chromatin

As a first step to characterising the function of MTBP in DNA replication, we analysed the chromatin recruitment of MTBP and Treslin at various times after adding sperm nuclei to interphase extract. MTBP and Treslin were recruited onto chromatin at 15 minutes, during the licensing phase and before replication initiation, when Cdc45 and PCNA were not yet present on chromatin (Fig. 3a). The MTBP and Treslin signal peaked at 30 minutes during S phase, when CMG proteins (Cdc45, Psf2 and Sld5), pre-IC proteins (TopBP1, RecQ4, Mcm10 and AND1/Ctf4) and DNA synthesis proteins (RPA, PCNA & the three replicative polymerases α, β and ε) were present on chromatin, and decreased during the later stages of S phase consistent with these (Figs. 3a and S5). In contrast, when sperm chromatin was incubated in metaphase-arrested extract, neither MTBP nor Treslin were recruited to the chromatin (Fig. 3b). These results indicate that MTBP and Treslin bind chromatin before S phase starts and peak during S phase, consistent with them being involved in replication initiation.

**Figure 3.**
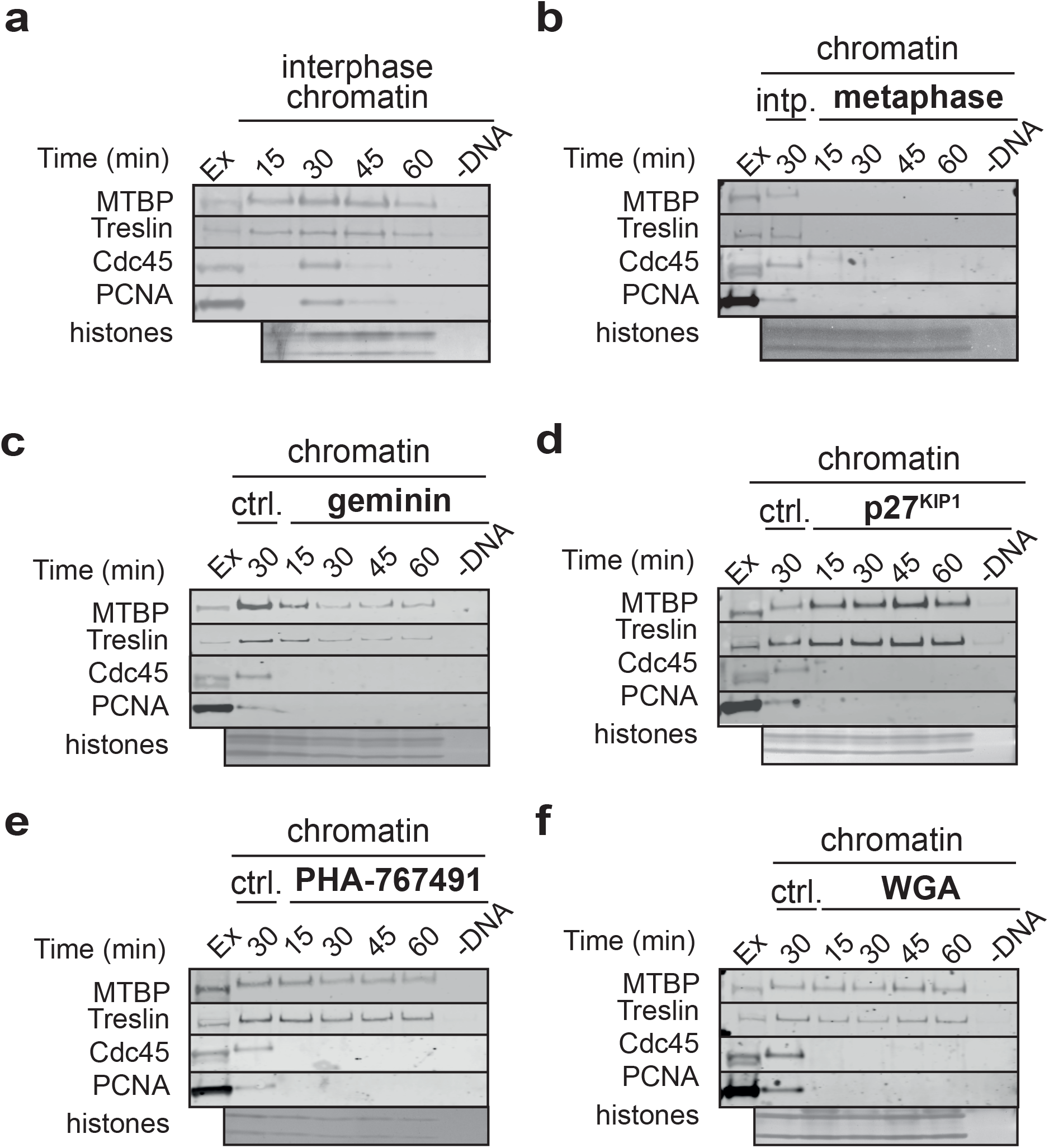
MTBP loading onto chromatin in metaphase and interphase extract. **a, b.** Sperm nuclei were incubated in interphase (a) or metaphase (b) extract Ex – extract; ctrl (b) – chromatin incubated for 30 min in interphase extract. **c-f.** Sperm nuclei were incubated in interphase extract for the indicated times in the presence of either (c) geminin or (d) p27^KIP1^, (e) PHA-767491 or (f) wheat germ agglutinin (WGA); chromatin was isolated, run on NuPage gels and immunoblotted for the indicated proteins. The lower portion of the gel was stained with Sypro Ruby to detect histones. Ex – extract; ctrl – chromatin incubated for 30 min in interphase extract without drug treatment.

We next carried out chromatin isolation from interphase extract supplemented with geminin to block replication licensing and prevent the MCM2-7 complex from loading onto chromatin(McGarry and Kirschner, 1998; Tada et al., 2001; Wohlschlegel et al., 2000). Figs. 3c and S6a show that whereas geminin significantly inhibited MTBP and Treslin recruitment onto chromatin, which peaked at 15 mins and then declined, all CMG helicase and all other pre-IC proteins are unable to associate with chromatin when licensing is inhibited. A similar decrease in MTBP and Treslin association with chromatin was observed in extract treated with RL5a, a small molecule which inhibits the ability of ORC to bind productively with DNA and which thereby inhibits origin licensing (Gardner et al., 2017) (Fig S6).

To verify whether MTBP requires S phase kinase activity to load onto chromatin, we inhibited the DDK and CDK activities that are required for the initiation of DNA replication in *Xenopus* egg extract (Blow and Nurse, 1990; Fang and Newport, 1991; Strausfeld et al., 1994; Jares and Blow, 2000; Walter, 2000). Treslin is a substrate for S phase CDKs, and its phosphorylation is required for it to bind to Dpb11/Cut5/TopBP1 at replication origins (Masumoto et al., 2002; Tanaka et al., 2007; Zegerman and Diffley, 2007; Kumagai et al., 2010; Natsume et al., 2013; Kumagai *et al*., 2011). DDK is recruited to MCM2-7 at licensed origins where it phosphorylates MCM2 and MCM4 (Jares and Blow, 2000; Walter, 2000; Poh *et al*., 2014; Alver *et al*., 2017).

Figs. 3d and S6b show that in extract treated with the CDK inhibitor p27^KIP1^, the Cdc45 and GINS (Psf2 and Sld5 proteins) components of CMG, pre-IC protein TopBP1 and the proteins required for DNA synthesis do not associate with chromatin, but MTBP and Treslin were loaded onto chromatin to higher levels compared to the control, consistent with a previous report that inhibition of CDKs causes a hyper-loading of Treslin in *Xenopus* egg extracts (Kumagai *et al*., 2010). This indicates that neither MTBP nor Treslin need S phase CDK activity to load onto chromatin, and that CDK activity actually reduces their loading. RecQ4 was also loaded onto chromatin at normal levels when CDK activity was inhibited (Fig S6).

When PHA-767491 was added to *Xenopus* extract to inhibit DDK activity (Montagnoli et al., 2008; Poh *et al*., 2014; Alver *et al*., 2017), MTBP and Treslin peaked on chromatin at 15 min at levels comparable to the control, but then declined, whereas other than the MCM proteins, all other components of CMG, all other pre-IC and the proteins required for DNA synthesis did not associate, (Fig. 3e and S5b). When we used Wheat Germ Agglutinin to inhibit nuclear pore function (Finlay et al., 1987) - thereby reducing the effective concentration of replication factors around DNA - Treslin-MTBP recruitment stayed at an approximately constant level comparable to the peak seen in control extract (Fig. 3f). This could be explained by the opposing effects of reducing the concentrations of Treslin-MTBP and DDK concentrations around DNA which would tend to reduce Treslin-MTBP loading, whilst at the same time reducing CDK activity around DNA which would tend to increase Treslin-MTBP loading.

The reduced chromatin association of Treslin-MTBP in extracts treated with either geminin or RL5a to inhibit licensing and PHA-767491 to inhibit DDK activity is in stark contrast to the hyperloading seen when CDK activity is inhibited by p27^KIP1^ (Figs. 3, S5 and S6). This removal of Treslin-MTBP from chromatin following peak association at 15 mins and hyperloading is consistent with nuclear formation and the concomitant increase of local CDK activity. To determine the combined effect of inhibiting both CDK activity and either licensing and DDK activity we isolated chromatin from egg extract, in increasing concentrations of salt, that had been treated either with p27^KIP1^, geminin or PHA-767491 alone or a combination of two inhibitors. Fig. S7 shows that either in the presence or absence of licensing or DDK activity, inhibition of CDK both increases and stabilises the association of Treslin-MTBP with chromatin, however it is only in the presence of both licensing and DDK activity that maximal binding is seen.

### Involvement of MTBP in DNA Replication

To verify the involvement of MTBP in DNA replication we immunodepleted MTBP from *Xenopus* egg extract using the MTBP-1 and MTBP-2 antibodies. The residual MTBP and Treslin in the depleted extracts was ∼5% (Fig. S8a). Extracts depleted with MTBP-1 or MTBP-2 antibodies showed a strongly reduced ability to support DNA replication (Fig. 4a). To verify that both the MTBP-1 and MTBP-2 antibodies depleted the same factor required for replication, we performed complementation assays, where equal volumes of depleted extract were mixed and then subjected to a replication assay. Mixtures of mock-depleted extract and MTBP-1 or MTBP-2 depleted extracts supported replication, whilst mixtures of MTBP-1 and MTBP-2 depleted extracts did not (Fig. S8b). This indicates that the MTBP-1 and MTBP-2 antibodies deplete the same essential factors required for DNA replication.

**Figure 4.**
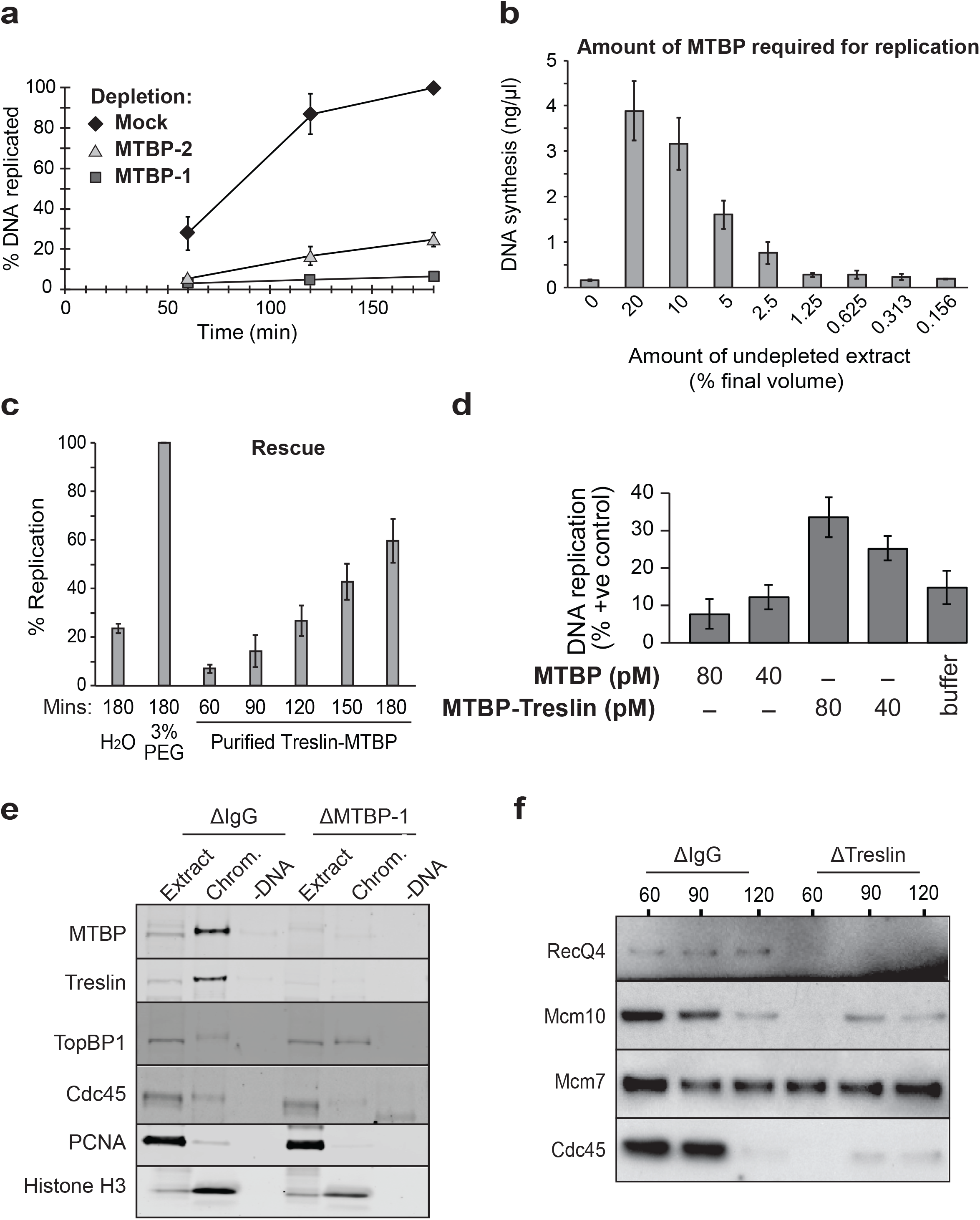
Immunodepletion of MTBP from *Xenopus* egg extract. **a.** DNA replication was assayed in the indicated depleted extracts (average of three independent experiments, error bars represent standard error). **b.** Undepleted extract was titrated into MTBP-1 depleted extract and DNA synthesis was assayed after three hours. Average of three independent experiments; error bars represent standard error. Sperm nuclei were added to a final DNA concentration of 3ng/µl. **c.** MTBP-1 depleted extract was incubated with H_2_O, 3% PEG or ammonium sulphate concentrated partially purified MTBP/Treslin complex diluted in H_2_O. Replication was assayed at the indicated times. Average of three independent experiments; error bars represent standard error. **d.** MTBP or Treslin–MTBP was delivered to MTBP-1 depleted extract at either 40 or 80 pM. Sperm chromatin was added and replication was assayed after three hours, expressed as a percentage of the replication achieved in depleted extract supplemented with an excess of the 3% PEG fraction. Average of three independent experiments; error bars represent standard error. **e.** Mock or MTBP-1 depleted extract were supplemented plus or minus sperm chromatin; after 90 mins chromatin was isolated, run on a NuPage gel and immunoblotted for the indicated proteins. **f.** Mock or Treslin-1 depleted extract were supplemented with sperm chromatin, chromatin was isolated at the indicated times, run on a NuPage gel and immunoblotted for the indicated proteins.

Even though in MTBP-depleted extracts DNA replication was significantly reduced the depleted extracts could still efficiently support complementary strand synthesis on single-stranded M13 plasmid DNA (Fig. S8c) and nuclear assembly was not impaired (Fig. S8d). To determine the amount of Treslin-MTBP complex required for DNA replication we performed serial dilutions of normal extract into MTBP-depleted extracts and tested the amount of DNA synthesis (Fig. 4b). This showed that 5% by volume of undepleted extract could rescue replication to an efficiency of ∼50% in the depleted extracts. This is equal to an MTBP concentration of ∼0.1 nM and a Treslin-MTBP tetramer concentration of ∼0.05 nM. Given the significantly longer S phase observed in immunodepleted extracts this is consistent with the hypothesis that MTBP is present at limiting levels for replication initiation.

We next carried out a rescue experiment adding an aliquot of partially-purified Treslin-MTBP (Fig. 2) to extract depleted with MTBP antibodies. Ammonium sulphate precipitation was used to concentrate the Treslin-MTBP sufficiently to allow assay, but residual ammonium sulphate in the sample was somewhat inhibitory to DNA replication. As positive control for rescue we used the maximum rescue achieved by Treslin-MTBP concentrated after precipitation with PEG (directly from extract Fig. 4c). The fraction containing the 600-fold enriched Treslin-MTBP complex rescued replication to 60% compared to the positive control. This highly efficient rescue strongly suggests that Treslin-MTBP is the replication factor missing in the depleted extracts.

We next asked whether MTBP alone is sufficient to rescue the replication phenotype of the MTBP-depleted extract, or whether the Treslin-MTBP complex is required. During anion exchange chromatography, ∼20% MTBP elutes before the Treslin peak, suggesting that MTBP is in slight excess in egg extract (Fig. 2). We compared the ability of fractions containing MTBP only or the Treslin-MTBP complex (Fig. 4d and S8e) to rescue DNA replication in MTBP-depleted extract. Whilst fractions containing Treslin-MTBP could rescue DNA replication to ∼30% of the positive control, comparable amounts of MTBP in the MTBP-only fraction were not able to rescue replication at all. These results suggest that MTBP alone is not sufficient to rescue the replication phenotype of the MTBP-depleted extract and that Treslin is an essential factor for replication.

To verify which step of DNA replication was impaired in the MTBP-depleted extracts, we carried out chromatin isolation from MTBP-depleted, Treslin-depleted or mock-depleted extracts 90 minutes after the addition of sperm DNA to the extracts. In the MTBP and Treslin-depleted extract TopBP1 was recruited to chromatin, but the recruitment of RecQ4, Cdc45 and PCNA was strongly reduced (Fig. 4e, f). Since maximal loading of Treslin-MTBP requires both licensing and DDK activity and in addition to these TopBP1 requires CDK activity we next determined the dynamic association of TopBP1 across the replication cycle in the absence of Treslin-MTBP. Fig 5a shows that TopBP1 chromatin association followed that of Treslin-MTBP, showing a peak of binding during S phase which then declines as S phase proceeds. In extract depleted of Treslin-MTBP, the association of TopBP1 with chromatin was reduced but did not decline at later times. When either licensing or CDK activity were inhibited by the addition of geminin or p27^KIP1^ the association of TopBP1 during peak S phase in the absence of Treslin-MTBP was maintained though at a reduced level (Figs. 5b and c). In combination these results suggest that although TopBP1 can associate with licensed chromatin in the presence of CDK activity, maximal binding during a normal S phase requires Treslin-MTBP. This extends previous reports of Treslin and MTBP depletion (Kumagai and Dunphy, 2017; Kumagai *et al*., 2010), and indicates that MTBP depletion impairs the assembly of the pre-IC required for the formation of the CMG helicase on chromatin.

**Figure 5.**
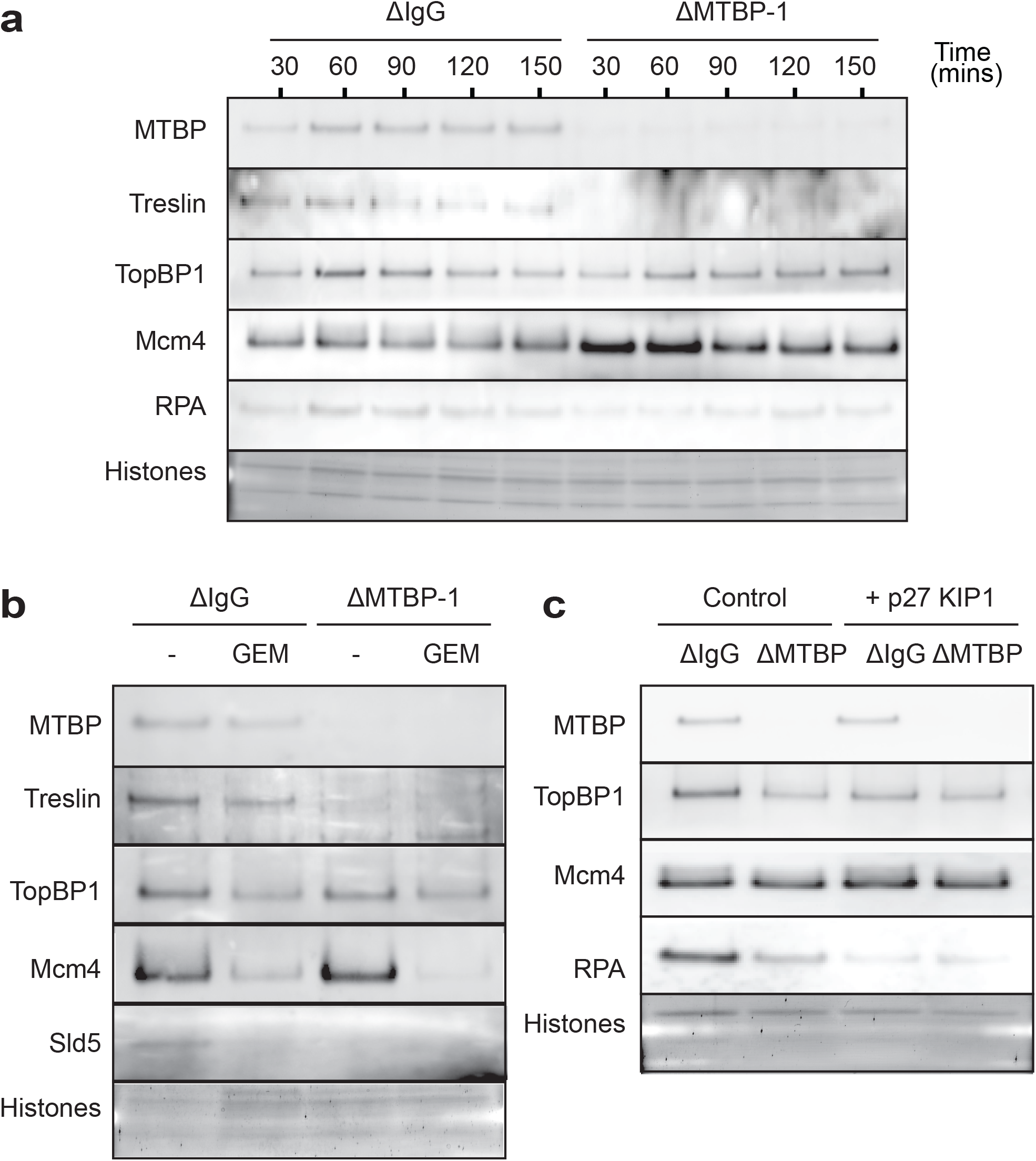
TopBP1 chromatin association in the absence of MTBP. **a**. Mock or MTBP-1 depleted extract was incubated with sperm chromatin for the indicated times, chromatin was isolated, run on a NuPage gel and immunoblotted for the indicated proteins. The lower portion of the gel stained with Sypro Ruby to detect histones. **b, c.** Mock or MTBP-1 depleted extract were supplemented with sperm chromatin and either ± (b) geminin or (c) p27^KIP1^; after 60 mins chromatin was isolated, run on a NuPage gel and immunoblotted for the indicated proteins. The lower portion of the gel stained with Sypro Ruby to detect histones.

To determine whether the Treslin-MTBP complex loaded onto DNA in the absence of CDK or DDK activity is functional, we carried out chromatin transfer experiments (Fig. 6a). Sperm chromatin was incubated for 60 min in extract supplemented with p27^KIP1^ or PHA-767491 to inhibit CDK or DDK activities; the chromatin was then isolated and transferred to MTBP-depleted extract to determine if the previously loaded MTBP and Treslin could support replication initiation. As control, the same DNA templates were added to extract mock-depleted with non-immune IgG. Chromatin from extracts treated with PHA-767491 and p27^KIP1^ were able to replicate efficiently in MTBP-depleted extract, indicating that the Treslin-MTBP complex loaded onto chromatin in these conditions is potentially functional (Fig. 6b).

**Figure 6.**
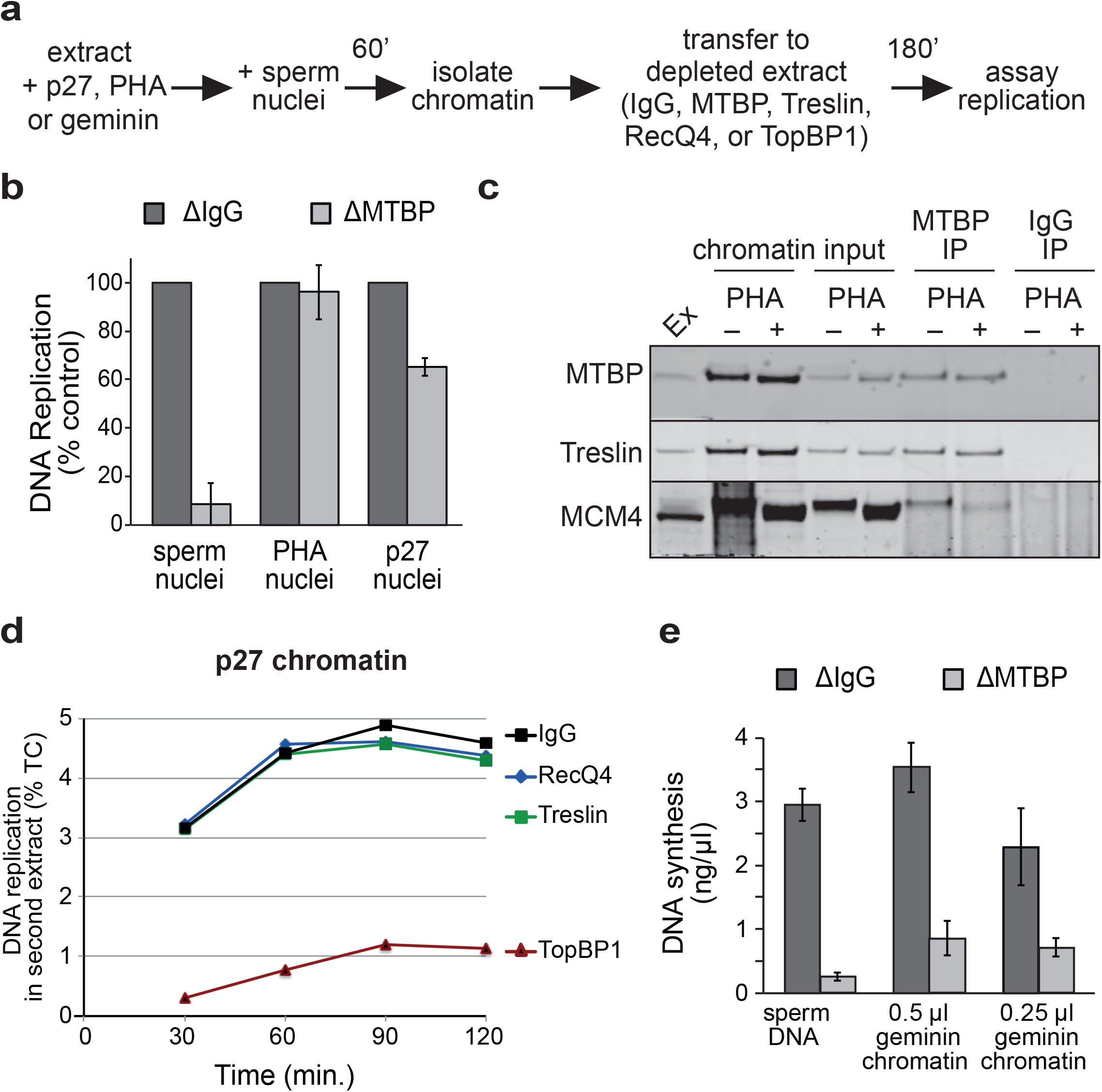
Treslin-MTBP complex regulation by replicative kinases. **a.** Schematic of nuclear transfer experiments **b.** Sperm nuclei were incubated for 60 mins in extract supplemented with p27^KIP1^ or PHA-767491; chromatin was isolated and transferred to extract immunodepleted with MTBP-1 antibodies or non-immune IgG; the total DNA synthesis after 180 mins was measured. Average of three independent experiments; error bars represent standard error. **c.** MTBP was immunoprecipitated from chromatin incubated in extract containing p27 plus or minus PHA-767491 and samples were immunoblotted for the indicated proteins. **d.** Sperm nuclei were incubated for 60 mins in extract supplemented with p27^KIP1^; chromatin was isolated and transferred to extract immunodepleted with Treslin-1, TopBP1, RecQ4 antibodies or non-immune IgG; the total DNA synthesis at the indicated times was measured. **e.** Sperm nuclei were incubated for 60 mins in extract supplemented with geminin; chromatin was isolated and transferred to extract immunodepleted with MTBP-1 antibodies or non-immune IgG; the total DNA synthesis after 180 mins was measured. Average of three independent experiments; error bars represent standard error.

To verify whether the interaction between Treslin-MTBP and MCM2-7 is strengthened by DDK activity, as occurs in yeast (Deegan *et al*., 2016), we treated interphase extract with p27^KIP1^ to block replication initiation and compared the amount of MCM co-precipitating with Treslin-MTBP plus or minus PHA-767491. Fig. 6c shows that the amount of MCM4 co-precipitating with Treslin-MTBP was lower when DDK activity was inhibited. Significantly, the residual MCM4 co-precipitating in the presence of PHA-767491 was hyperphosphorylated compared to bulk chromatin, consistent with the idea that Treslin-MTBP binds more strongly to MCM2-7 that has been phosphorylated by DDK. Consistent with the ability of Treslin-MTBP and RecQ4 but not TopBP1 to associate with chromatin when CDK activity is inhibited, chromatin isolated from p27^KIP1^ treated extract could replicate efficiently in Treslin-MTBP and RecQ4 but not TopBP1 depleted extracts (Fig 6d).

In order to determine whether the low quantity of Treslin-MTBP recruited to chromatin in the absence of licensing was functional, we isolated chromatin from extracts treated with geminin, which contained ∼20% of control levels of Treslin-MTBP, and assayed its ability to replicate in MTBP-depleted extract. Fig. 6e shows that the chromatin from geminin-treated extract replicated to <20% of control levels in MTBP-depleted extract. This suggests that origin licensing and MCM2-7 loading are required for Treslin-MTBP to be efficiently recruited to chromatin in a functional manner, but that the reduced amount of Treslin-MTBP loaded onto DNA in the absence of MCM2-7 is present in a potentially functional form.

### DDK Activity Regulates Treslin-MTBP-TopBP1 complex formation

We have shown that both licensing and DDK activity are required both to increase and strengthen the association of Treslin-MTBP with chromatin. This is consistent with the hyperphosporylation of Mcm4 that we have previously shown correlates with DNA replication (Poh *et al*., 2014; Alver *et al*., 2017). The hyperphosphorylation of Mcm4 is reversed by Protein Phosphatase 1 (PP1) targeted to chromatin by Rif1. However, chromatin with hyperphosphorylated Mcm4 isolated from egg extract treated in the presence of I-2 an inhibitor of PP1 does not replicate fully when transferred to an extract in which both DDK and PP1 activity have been inhibited (Poh *et al*., 2014). This suggests either that some dephosphorylation of Mcm4 occurs even in the presence of I-2 or that there is a second DDK substrate required for the efficient initiation of DNA replication. In considering possible additional DDK substrates, we noted that chromatin-associated Treslin migrated slower in gels compared to the band present in extract, and that this slower migration was reduced when DDK activity was inhibited (Fig. 6c), suggesting that Treslin itself maybe a DDK substrate. We therefore further characterised the change in Treslin migration in the presence of DDK inhibition by PHA-767491. We treated egg extract with inhibitors of either DDK or CDK activity alone or in combination, isolated chromatin following nuclear assembly during S phase and analysed Treslin mobility by immunoblotting. Fig. 7a shows that upon inhibition of either DDK or CDK activity alone Treslin migrates faster than seen in control but to a different extent; in combination, inhibition of both DDK and CDK activity, results in Treslin migrating faster than in each of the treatments alone suggesting that both DDK and CDK independently contribute to the changes in Treslin mobility and that Treslin is a target of DDK activity.

**Figure 7.**
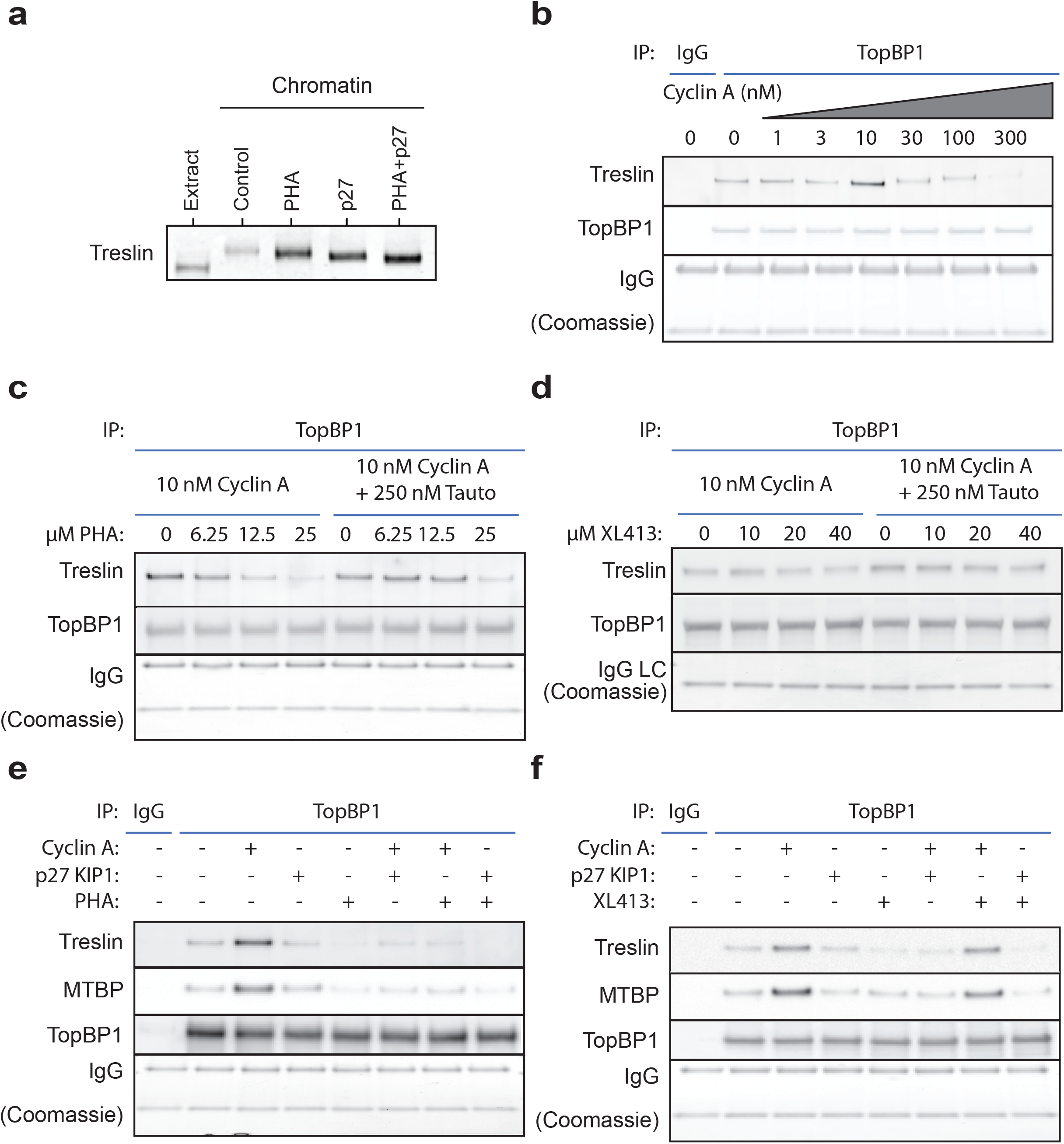
DDK & PP1 regulate CDK-mediated Treslin-MTBP & TopBP1 interaction. **a.** Sperm nuclei were incubated in extract supplemented with either p27^KIP1^ or PHA-767491 alone or in combination, as indicated, for 60 mins; chromatin was isolated, run on a NuPage gel and immunoblotted for Treslin. **b-e.** Diluted and clarified egg extract was supplemented with cyclin A, p27^KIP1^, PHA-767491, XL413, Tautomycetin, either alone or in combination, as indicated and incubated for 15 mins. Samples were immunoprecipitated using either control IgG or anti-TopBP1-antibody bound protein-G Dynabeads. Washed precipitates were subjected to SDS-PAGE on a NuPage gel. The upper portion of the gel was immunoblotted for Treslin, MTBP and TopBP1 as indicated and the lower portion of the gel stained with Coomassie to visualise the Heavy and Light chains of IgG.

Treslin-MTBP associates with chromatin bound MCM2-7 and also forms a tripartite complex with TopBP1, thereby coupling replication licensing and pre-IC formation. We reasoned that since Treslin-MTBP associates more strongly with DDK-mediated hyperphosphoryated Mcm4 at licensed replication origins that DDK activity may also be required in addition to CDK activity to mediate Treslin-MTBP and TopBP1 interaction. We first determined the optimal level of CDK activity required to promote interaction between Treslin-MTBP and TopBP1. In the absence of nuclear formation CDK activity in egg extract is low; upon nuclear formation and protein import the local concentration of CDK activity within nuclei increases at least 25 fold, a level sufficient to induce replication initiation. In egg extract the CDK1 catalytic subunit is in ∼50 fold excess compared to its activating cyclin partners so addition of cyclin stimulates CDK activity *in vitro* (Strausfeld *et al*., 1994). To mimic the nuclear environment in which Treslin-MTBP-TopBP1 interaction occurs we titrated cyclin A into 5-fold diluted and clarified egg extract in the absence of DNA and assessed the formation of the Treslin-MTBP-TopBP1 complex by determining the recovery of Treslin in TopBP1 immunoprecipitates (Fig. 7b). The Treslin-TopBP1 interaction was maximal upon addition of 10 nM cyclin A into egg extract but declined thereafter as cyclin A concentration is further increased, concentrations at which the little Treslin that is seen to interact with TopBP1 is increasingly retarded in mobility. Maximal Treslin-TopBP1 interaction upon the addition of 10 nM cyclin A is consistent with the level of Cdk2-cyclin A required to support maximal DNA replication in egg extracts, above which the extract enters a mitotic-like state (Strausfeld *et al*., 1994).

We next determined the effect of the addition of inhibitors of DDK to egg extract on cyclin A-stimulated Treslin-TopBP1 interaction. Figs. 7c and 7d show that addition of the DDK inhibitors PHA-767491 or XL413 reduced the interaction of Treslin with TopBP1. Compared to PHA-767491, XL413 is a more specific but less potent inhibitor of DDK-mediated chromatin associated Mcm4 phosphorylation in egg extract (Alver *et al*., 2017) and as expected, addition of XL413 to egg extract inhibited Treslin-TopBP1 interaction to a lesser extent. The inhibition of Treslin-TopBP1 complex formation by inhibition of DDK was reversed upon co-addition of the PP1 inhibitor Tautomycetin into extract, consistent with the role of PP1 in reversing DDK-mediated phosphorylation of Mcm4 (Alver *et al*., 2017; Poh *et al*., 2014). These results suggest that DDK activity is required for the interaction of Treslin-MTBP and TopBP1 and that this is subject to regulation by PP1.

Whereas CDK activity in the absence of nuclear formation is low in egg extract, DDK activity is detectable but reduced (Alver *et al*., 2017; Poh *et al*., 2014). To further address the role of DDK activity on formation of the Treslin-MTBP-TopBP1 complex we next determined the effect of combined inhibition of both DDK and CDK activity on their interaction. Figs. 7e and f show that cyclin A’s ability to stimulate formation of the Treslin-MTBP-TopBP1 complex is curbed both by inhibiting CDK activity with p27^KIP1^ and by inhibiting DDK activity with either PHA-767491 or XL413. Inhibition of both CDK and DDK activities reduced complex formation to the greatest extent but no more so than with PHA-767491 alone. This suggests both that the formation of the Treslin-MTBP-TopBP1 complex is dynamic and that DDK activity is required for the CDK-dependent formation of the complex.

The dynamic nature of the Treslin-MTBP-TopBP1 interaction is consistent with the role we identified for PP1 in reversing DDK inhibition of complex formation (Figs. 7c and d). Maximal interaction required both optimal CDK (Fig. 7b) and DDK activity. Figs. 8a and b show that addition of the PP1 inhibitor Tautomycetin alone stimulated the interaction between Treslin-MTBP and TopBP1, although not to the same extent as the addition of cyclin A. Maximal complex formation was seen when both DDK and CDK activity were stimulated by the combined addition of both Tautomycetin and cyclin A. Whereas addition of either PHA-767491 or XL413 reversed the stimulation seen upon addition of Tautomycetin, the further addition of cyclin A could overcome this to some extent. These results are consistent with the idea that DDK activity is actually more important than CDK activity for the formation of the Treslin-MTBP-TopBP1 complex. Complex formation was more strongly reduced by DDK inhibition than by CDK inhibition (Fig 6e, f). Stimulation of DDK-dependent phosphorylation with tautomycetin promoted complex formation in the absence of cyclin A, but only very low levels of complex formation occurred when DDK activity was inhibited in the presence of cyclin A. Taken together these results suggest that DDK activity is essential for the formation of the Treslin-MTBP-TopBP1 complex, whilst optimal CDK activity further strengthens the interactions.

**Figure 8.**
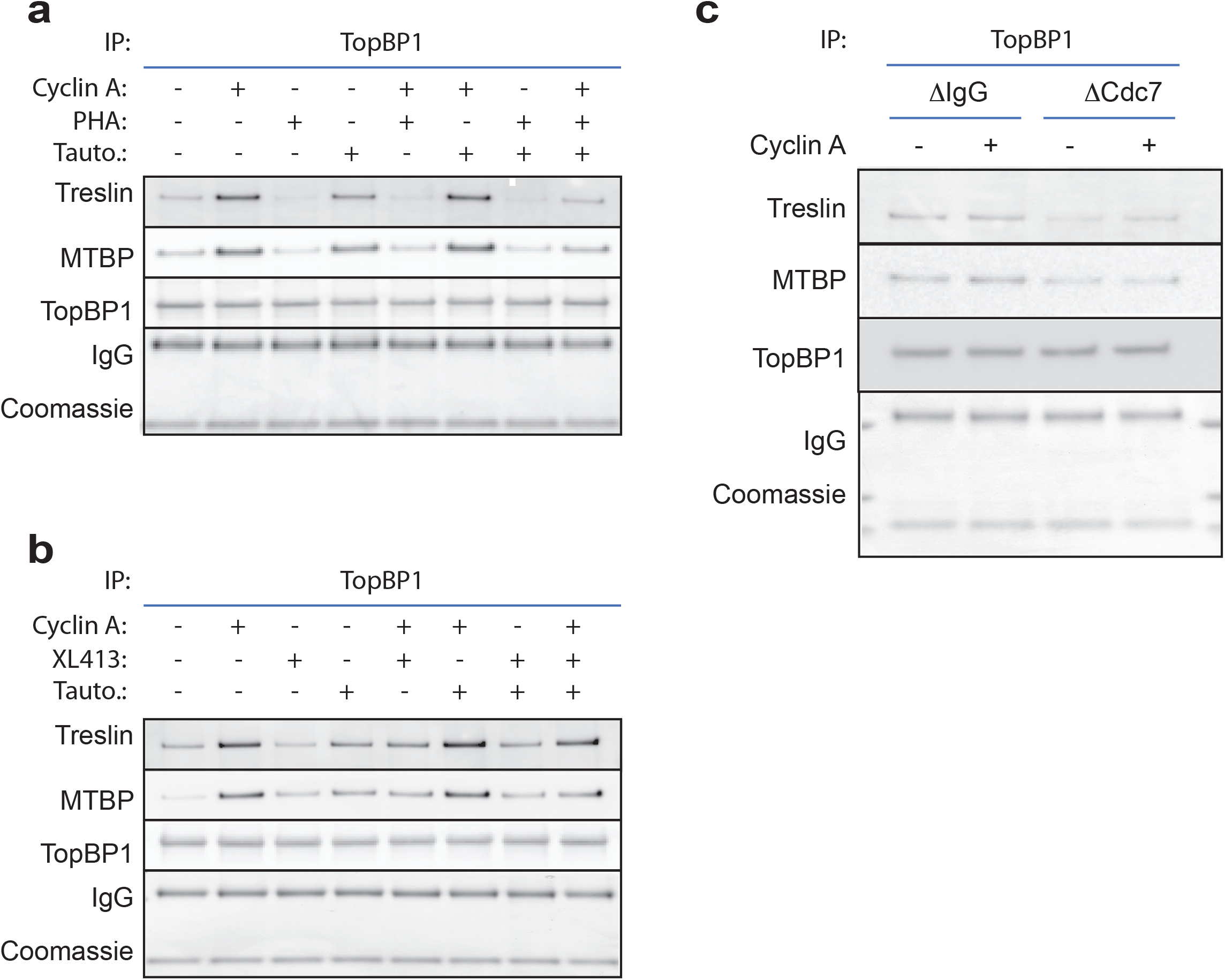
PP1 reverses DDK mediated inhibition of CDK-mediated Treslin-MTBP & TopBP1 complex formation. **a, b.** Diluted and clarified egg extract was supplemented with cyclin A, p27^KIP1^, PHA-767491, XL413, Tautomycetin, either alone or in combination, as indicated and incubated for 15 mins. Samples were immunoprecipitated using either control IgG or anti-TopBP1-antibody bound protein-G Dynabeads. Washed precipitates were subjected to SDS-PAGE on a NuPage gel. The upper portion of the gel was immunoblotted for Treslin, MTBP and TopBP1 as indicated and the lower portion of the gel stained with Coomassie to visualise the Heavy and Light chains of IgG. **c.** <u>Extract immunodepleted with a control IgG or anti-Cdc7 (DDK) antibody was either supplemented or not with 10 nM cyclin A, as indicated and incubated for 15 mins.</u> Samples were immunoprecipitated using either control IgG or anti-TopBP1-antibody bound protein-G Dynabeads. Washed precipitates were subjected to SDS-PAGE on a NuPage gel. The upper portion of the gel was immunoblotted for Treslin, MTBP and TopBP1 as indicated and the lower portion of the gel stained with Coomassie to visualise the Heavy and Light chains of IgG.

To address directly the role of DDK activity in Treslin-MTBP-TopBP1 complex formation we next prepared extract immunodepleted of Cdc7. Immunodepletion of at least 95% of Cdc7 from the extract left MTBP levels similar to those in control depleted extract (Fig. S9a). Consistent with the results presented above using both PHA-767491 and XL413 to inhibit DDK activity, removal of Cdc7 from egg extract reduced the interaction of Treslin-MTBP with TopBP1 both in the presence and absence of additional cyclin A (Figs 8c and S9b).

## Discussion

In this study we show that in *Xenopus* egg extracts, MTBP and Treslin form an elongated complex that is recruited to DNA before replication initiates. The Treslin-MTBP complex can be recruited to DNA in a functional form in the absence of replication licensing, DDK or CDK activity, but licensing and DDK activity stabilizes the interaction between Treslin-MTBP and chromatin bound MCM2-7. We find that the Treslin-MTBP complex is present in extract at a low concentration expected to be rate-limiting for replication initiation. We also demonstrate an important and previously unrecognised role for DDK activity in promoting the interaction of Treslin-MTBP with TopBP1. Since DDK also has an essential role in phosphorylating chromatin-bound MCM2-7, this suggests that DDK plays a central role in directing the recruitment of Treslin-MTBP and TopBP1 to licensed replication origins, thereby selecting which origins will subsequently initiate.

Treslin and MTBP co-immunoprecipitate with each other in both metaphase and interphase extract indicating that they interact throughout the cell cycle, as previously reported in human cells and *Xenopus*, and similar to Sld3 and Sld7 in budding yeast (Tanaka *et al*., 2011b; Boos *et al*., 2013; Kumagai and Dunphy, 2017). Using a combination of sucrose gradient sedimentation and gel filtration we show that the Treslin-MTBP complex has a native molecular weight of 626 KDa, close to (± ≤1%) the value expected for a tetramer composed of two molecules of MTBP (96 KDa) and two molecules of Treslin (220 kDa), that is 632 kDa. The distinct behaviours of *Xenopus* Treslin-MTBP on gel filtration and sucrose gradients suggests that it forms a highly elongated complex. This result is in line with the model proposed for Sld3 and Sld7 in yeast - obtained from studies of the crystal structures of domains of Sld3 and Sld7 - which proposed that two molecules of Sld7 could bridge two molecules of Sld3, with one loaded onto each of the hexamers of MCM2-7 loaded at an origin to allow bi-directional initiation (Itou *et al*., 2015).

MTBP and Treslin also interact on chromatin during S phase and at this time they also interact with Cdc45 and the GINS complex, consistent with the idea that they function during replication initiation. Immunodepletion of MTBP from *Xenopus* egg extract co-depleted Treslin with impairment of DNA replication due to lack of CMG assembly, supporting the idea that Treslin-MTBP is required for CMG formation. The capacity to initiate replication could be restored to MTBP-depleted extract by co-addition of a partially-purified Treslin-MTBP complex, but not by fractions containing MTBP alone. This is consistent with a recent report which shows that DNA replication in an MTBP-depleted extract can be rescued with a combination of recombinant Treslin and MTBP, but not either alone(Kumagai and Dunphy, 2017).

We found that MTBP was recruited onto chromatin during licensing and peaked in S phase, similar to Treslin. The chromatin recruitment of MTBP and Treslin occurred in the absence of licensing, DDK, CDK or nuclear assembly (Fig 9a i), but did not occur in metaphase arrested extract. Chromatin transfer experiments showed that the Treslin-MTBP recruited onto chromatin in the absence of licensing, DDK or CDK could efficiently support replication in MTBP-depleted extract; this suggests that Treslin-MTBP binds unlicensed chromatin in a form that can be efficiently activated when other initiation activities are supplied. These results differ from studies in yeast, where the loading of Sld3 and Sld7 onto origins depends on licensing and DDK activity (Yeeles *et al*., 2015). However, we show that in extract where licensing was abolished by addition of active geminin or RL5a, the loading of MTBP and Treslin onto chromatin was strongly reduced compared to control, indicating that the MTBP and Treslin loading that occurs before licensing is relatively unstable (Fig 9a ii). MCM2-7 is known to be phosphorylated by DDK and this phosphorylation is required for pre-IC formation (Jares and Blow, 2000; Walter, 2000; Poh *et al*., 2014; Alver *et al*., 2017; Sheu and Stillman, 2006). We show here, that consistent with data in yeast (Deegan *et al*., 2016) the physical interaction between the Treslin-MTBP and MCM2-7 was strengthened by DDK activity (Fig 9a iii). The Treslin-MTBP bound to chromatin in the absence of either licensing or DDK and CDK activity can support DNA replication suggesting that it is in a potentially active configuration.

**Figure 9.**
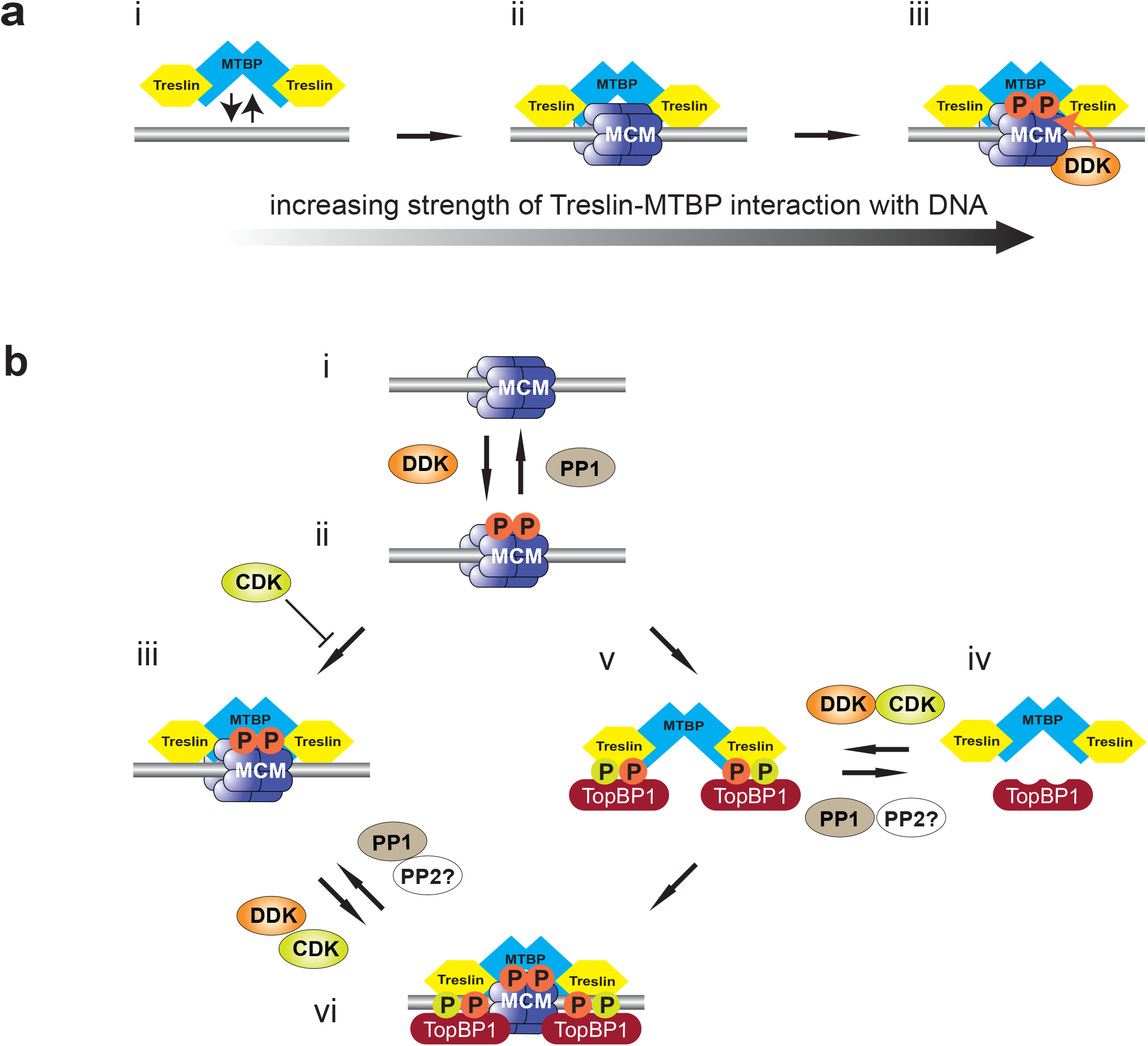
Model for the role of Treslin-MTBP, DDK and CDK in pre-IC formation. **a.** Model of the interaction of Treslin-MTBP with replication origins. i) Treslin-MTBP can bind weakly to DNA. ii) Stronger binding of Treslin-MTBP to licensed origins (DNA-bound Mcm2-7 double hexamers). iii) Strongest binding of Treslin-MTBP to licensed origins where Mcm2-7 has been phosphorylated by DDK. **b.** Model for pre-IC formation. i) Licensed origin (DNA-bound Mcm2-7 double hexamers). ii) DNA-bound Mcm2-7 is phosphorylated by DDK. Phosphorylation is reversed by PP1 recruited to origins by Rif1. iii) Treslin-MTBP is recruited to DDK-phosphorylated Mcm2-7. This interaction is opposed by CDK, most likely due to CDK phosphorylation of Treslin. iv) Soluble Treslin bound to MTBP can be phosphorylated by both CDK and DDK. v) Phosphorylated Treslin-MTBP binds to TopBP1. vi) The Treslin-MTBP-TopBP1 complex is recruited to DDK-phosphorylated Mcm2-7 to form the pre-IC.

Whereas both licensing and DDK activity are required to both increase and strengthen the interaction of Treslin-MTBP with chromatin, CDK activity reduces this interaction (Fig 9B i-iii). Treslin is a CDK substrate and interacts with TopBP1 through its BRCT motifs following CDK-mediated phosphorylation of Treslin S1001 (Kumagai *et al*., 2010; 2011; Boos et al., 2011). It is therefore likely that CDK-dependent phosphorylation of Treslin reduces its affinity for MCM2-7 whilst increasing its affinity for TopBP1. Therefore, as CDK activity rises during S phase, Treslin-MTBP is likely to be recruited to replication origins as part of a complex with TopBP1 (Fig 9b iv-v) and that this is a precursor to formation of the pre-IC (Fig 9B vi). We find that in addition to the known regulation of the Treslin-TopBP1 interaction by CDK that DDK activity also plays a major role (Fig 9b v-vi). Inhibition of DDK activity by either addition of PHA-767491 or XL413, or depletion of Cdc7 restricts the interaction between Treslin-MTBP and TopBP1, even in the presence of an optimal concentration of cyclin A to support complex formation, identifying the DDK dependence of CDK-mediated complex formation. In addition, we find that the DDK-mediated regulation of complex formation is subject to reversal by PP1. By increasing and decreasing DDK and CDK activity separately, we provide data suggesting that that DDK activity is essential for the formation of the Treslin-MTBP-TopBP1 complex, whilst optimal CDK activity further strengthens the interactions.

Given that Treslin-MTBP binds to chromatin prior to nuclear assembly when DDK activity is detectable but reduced and before CDK activity rises to support replication initiation, it may be that TopBP1 is initially directed to Treslin-MTBP associated with licensed and DDK phosphorylated origins, forming a complex directly at the origin (Fig 9b i-iii), but that as CDK activity rises during S phase the Treslin-MTBP-TopBP1 complex plays a major role in driving formation of the pre-IC (Fig 9B vi-vi).This suggests that in addition to ensuring Treslin-MTBP-TopBP1 complex formation, CDK activity facilitates the removal of Treslin-MTBP from unlicensed chromatin, thus ensuring redistribution of the complex to only licensed and DDK-phosphorylated origins to maximise the levels of origin firing. DDK activity thus co-ordinates the coupling of replication licensing to pre-IC formation and therefore, ultimately, CMG formation, by both directing the correct chromatin binding of Treslin-MTBP and mediating its interaction with TopBP1.

Studies in *S. cerevisiae* suggest that Sld3, Sld2, Dpb11 and Dbf4 are present at limiting concentrations (Mantiero et al., 2011; Tanaka *et al*., 2011b; Tanaka *et al*., 2011a). Moreover, overexpression of these factors together with overexpression of Sld7 and Cdc45 increased the efficiency of origin firing, including the firing of normally dormant origins (Mantiero *et al*., 2011). In *Xenopus* embryos it has been found that Treslin, TopBP1, Drf1 and RecQ4 are limiting for replication after the mid blastula transition and the levels of these factors regulates the speed of cell division (Collart et al., 2013). We quantified the amount of MTBP in *Xenopus* egg extract and found that the concentration of the Treslin-MTBP tetramer is ∼1 nM. This is significantly lower than most other replication proteins and is lower than the estimated 4 nM of replication origins that can be efficiently replicated in *Xenopus* embryos and egg extracts (Blow and Laskey, 1986; Blow *et al*., 2001). Consistent with this, we showed that the replication rate in extract is highly sensitive to the concentration of Treslin-MTBP. This is also consistent with a previous report showing that the majority of Treslin becomes chromatin-bound as embryos approach the Mid-Blastula Transition (Collart *et al*., 2013). The low concentration of Treslin-MTBP in embryos is therefore likely to be rate-limiting for replication initiation as embryos approach the Mid-Blastula Transition and DNA concentration approaches the maximum capacity observed in egg extracts.

In yeast, Sld3 is required for replication initiation but is not part of the replisome and is not required for replication fork progression (Gambus et al., 2006; Kanemaki and Labib, 2006). The chromatin binding of *Xenopus* Treslin-MTBP supports this idea, as Treslin-MTBP is lost from chromatin in the later part of S phase when some replisomes are still active. As an initiation activity, Treslin-MTBP, possibly also associated with TopBP1, is therefore expected to be released from origins as they initiate, potentially being able to bind again to unfired origins to promote their initiation. At the maximum physiological concentration of DNA in *Xenopus* egg extract (close to the DNA concentration at the Mid-Blastula Transition), the concentration of active origins is ∼4 nM (Blow and Laskey, 1986; Blow *et al*., 2001). We estimate ∼1 nM of the Treslin-MTBP tetramer in egg extract, meaning that each molecule would have to drive the initiation of an average of 4 origins at this maximum DNA concentration. This is consistent with previous work on the kinetics of origin firing in *Xenopus* extract which estimated that S phase is driven by ∼3 cycles of origin firing (Luciani et al., 2004). Furthermore, given the known role of the inhibition of DDK activity in checkpoint signalling following the interruption of the progression of DNA replication, the identification of DDK and PP1 as regulators of Treslin-TopBP1 complex formation provides a novel point at which DNA replication could be regulated following checkpoint activation.

Not all licensed replication origins initiate in any given S phase, with the majority remaining dormant but able to initiate in case of replication stresses (Woodward et al., 2006; Ge et al., 2007; Ibarra et al., 2008; Blow and Ge, 2009; Ge and Blow, 2010). In *Xenopus* egg extracts, <10% of DNA-bound Mcm2-7 initiate in an undisturbed S phase (Mahbubani et al., 1997; Edwards et al., 2002; Woodward *et al*., 2006), meaning that at Mid-Blastula Transition DNA concentrations, the concentration of DNA-bound Mcm2-7 is at least 50 nM. Given the data we present here, it is highly likely that Treslin-MTBP plays a crucial role in determining which of the licensed origins are selected for initiation. In egg extract, there is high DDK activity so that almost all MCM2-7 loaded onto DNA become phosphorylated by DDKs (Poh *et al*., 2014; Alver *et al*., 2017). The loading of the limited amounts of Treslin-MTBP is the earliest known event in the initiation process that selects the origins that will go on to initiate, as the other limiting factors (TopBP1 and RecQ4) act after Treslin-MTBP has been loaded. The correct selection of replication origins is important for maintaining genome stability by regulating the distribution of active replication forks in response to replication stresses (Ge *et al*., 2007; Blow and Ge, 2009; Ge and Blow, 2010). We therefore predict that over- or under-expression of Treslin-MTBP causes defects in correct origin selection, providing an explanation for why MTBP is misregulated in certain cancers(Grieb *et al*., 2014a).

## Significance

Prior to S phase onset, future origins of DNA replication are ‘licensed’ for use by being encircled by inactive double hexamers of Mcm2-7. During S phase the combined action of the two S phase promoting kinases, Dbf4-dependent kinase (DDK) and cyclin-dependent kinase (CDK), drive the association of Mcm2-7 with Cdc45 and the GINS protein complex to form the active Cdc45-MCM-GINS (CMG) replicative helicase. This activation requires the function of several additional pre-Initiation Complex proteins of which Treslin-MTBP is one. Using the *Xenopus* egg cell-free system we describe here a detailed biochemical characterisation of MTBP complex formation with Treslin, Treslin-MTBP’s role in DNA replication initiation and its regulation by both DDK and CDK. We find that Treslin and MTBP interact across the cell cycle and consistent with their role in DNA replication initiation maintain association on replicating chromatin during S phase. We show that the strong association of Treslin-MTBP with chromatin requires both Mcm2-7 and DDK activity and that both DDK and CDK activity have a role in ensuring correct Treslin-MTBP chromatin association. Most interestingly, however, we show for the first time that the DDK dependent interaction between Treslin-MTBP and TopBP1, another key pre-Initiation Complex protein. This identifies DDK and Treslin-MTBP as key co-ordinators of replication licensing and helicase activation.

## Author Contributions

IV produced the anti-MTBP antibodies with PJG, performed characterisation of Treslin-MTBP complex formation and purified Treslin-MTBP, performed chromatin isolation experiments and developed and characterised the MTBP depletion. PJG identified MTBP, produced the anti-MTBP antibodies with IV, developed the Treslin-MTBP purification with IV, performed chromatin isolation and IP experiments for TopBP1 studies and provided oversight of the experimental work. GSC produced and characterised the anti-TopBP1 and anti-Treslin antibodies and performed chromatin isolation experiments. JJB led the project and co-wrote the paper with PJG and IV.

## Acknowledgements

This work was supported by CR-UK programme grant C303/A14301 and Wellcome Trust Investigator award WT096598MA. I.V. was supported by a BBSRC studentship. We would like to acknowledge help from Dr. Sara ten Have and Kelly Hodge of the GRE Proteomics Support Team at the University of Dundee (Wellcome Trust Strategic Award 097945/Z/11/Z). The authors declare that they have no conflicts of interest with the contents of this article.

## STAR Methods

### CONTACT FOR REAGENT AND RESOURCE SHARING

Requests for resources, reagents and further information should be directed to and will be fulfilled by, the Lead Contact, J. Julian Blow.

### EXPERIMENTAL MODEL AND SUBJECT DETAILS

#### Xenopus laevis

Wild type, sexually mature (≥ 1 year old) female and male South African Clawed frogs (Xenopus laevis) born and reared in the UK (University of Plymouth) were used in this study for the production of unfertilised eggs (from which extracts were prepared) and sperm, respectively. Frogs were maintained at 19°C in particulate filtered, dechlorinated water, at a density of ≤15 animals per 60 l tank, in a purpose built ‘aquacentre’ and were maintained by a professional staff at the University of Dundee adhering to Home Office (UK Government) animal husbandry guidelines; the animals have access to a Home Office (UK Government) approved veterinary surgeon. The frogs were fed a vegetable and fish based diet (Aquatic Diets 3, Mazuri Zoo Foods) 2-3 times per week, as required.

#### Experimental procedures

##### Recombinant Antigens, Antibodies and Inhibitors

Polyclonal antibodies were raised in rabbit against the following polypeptides: MTBP-2: 125 N-terminal amino acids of MTBP with 6-histidine tag at C- terminus (MERFVLCIHWERRAEQQQPVPQGLVYAQDIYTQLKEYSTNCTSTFPACSLTGNPGIRKWFFALQSLYGFSQFCSSDWEDLCPAVTTDDSEEPVQTALDECLDALQFPDGEDDNSRDSISQTNLFE) MTBP-1: 103 C-terminal amino acids of MTBP with 6-histidine tag at N-terminus (APSVPAQSKPSSHLELEQKESRSQKHNRMLKEVVSKTLQKHSIGVEHPCYAACNQRLFEISKF FLKDLKTSRGLL DEMKKAASNNAKQVIQWELDKLKKK). Antibodies were purified using rProtein-A Sepharose fast flow beads (GE Healthcare) or affinity purified using HiTrap NHS-Activated HP columns (GE Healthcare). For Treslin, polyclonal antibodies were raised in two different rabbits against the N-terminal 487 amino acids of *Xenopus* Treslin, with an N-terminal 6-histidine tag. For TopBP1, polyclonal antibodies were raised in sheep against the N-terminal 596 amino acids of *Xenopus* TopBP1, with an N-terminal 6-histidine tag.

Commercial antibodies used in this study are: anti-histone H3 FL-136 rabbit (Santa Cruz sc-10809); anti-hMTBP (MTBP-S; Sigma-Aldrich HPA025694); anti-hPCNA (PC10) (Santa Cruz sc-56). Antibodies against *Xenopus* proteins: Cdc45 and Sld5 (Gambus et al., 2011); MCM4(Prokhorova and Blow, 2000); ORC1 (Rowles et al., 1996); ORC2 (Oehlmann et al., 2004); TOP2A (Hirano and Mitchison, 1993). Anti–*Xenopus* WRN rabbit polyclonal antibody was a gift from H. Yan (Fox Chase Cancer Center, Philadelphia, USA) (Yan et al., 2005). Anti-rabbit IgG, anti-sheep IgG and anti-mouse–HRP linked antibodies were from New England Biolabs. Anti-rabbit IgG, anti-sheep IgG and anti-mouse IgG–Dylight 800 or Dylight 680 linked antibodies were from Thermo Fisher Scientific.

Recombinant geminin^DEL^ was as described (Thomson et al., 2010). p27^KIP1^ was described in (Poh *et al*., 2014). PHA-767491 was produced by Division of Signal Transduction Therapy, University of Dundee. *Triticum vulgaris* Wheat Germ Agglutinin was purchased from Sigma-Aldrich (L9640).

#### Xenopus egg extract methods

Metaphase-arrested *Xenopus laevis* egg extract and demembranated *Xenopus* sperm nuclei were prepared as described (Gillespie et al., 2012; Gillespie *et al*., 2016). Female frogs were primed with 50 units of Folligon (Pregnant Mare Serum Gonadotrophin) 3 days before the eggs were required to increase the number of stage 6 mature oocytes and 2 days later, were injected with 500 units Chorulon (Chorionic Gonadotrophin) to induce ovulation. Frogs were placed in individual laying tanks at 18-21°C in 2 l 1x MMR egg laying buffer, prepared from a 10x stock (1 M NaCl, 20 mM KCl, 10 mM MgCl_2_, 20 mM CaCl_2_,1 mM EDTA, 50 mM HEPES-NaOH, pH 7.8). The following morning, eggs were collected and rinsed in 1x MMR to remove any non-egg debris. Washed eggs were dejellied in 2% w/v cysteine (pH 7.8), washed in XBE2 (1x XB salts, 1.71% w:v sucrose, 5 mM K-EGTA, 10 mM HEPES-KOH, pH 7.7; 10x XB salts: 2 M KCl, 40 mM MgCl_2_, 2 mM CaCl_2_) and then into XBE2 containing 10 µg/ml leupeptin, pepstatin and aprotinin. Dejellied and washed eggs were centrifuged in 14 ml tubes, containing 1 ml XBE2 plus protease inhibitors containing 100 µg/ml cytochalasin D, at 1400 x g in a swinging bucket rotor for 1 min at 16°C to pack the eggs, after which excess buffer and dead eggs were removed. Packed eggs were crushed by centrifugation at 16,000 x g in a swinging bucket rotor for 10 min at 16°C. The dirty brown cytoplasmic layer was collected using a 20G needle and a 1 ml syringe via side puncture. From this point onwards the extract was kept on ice. The crude extract was supplemented with cytochalasin D, leupeptin, pepstatin and aprotinin all to a final concentration of 10 µg/ml, 1:80 dilution of energy regenerator (1 M phosphocreatine disodium salt, 600 µg/ml creatine phosphokinase in 10 mM HEPES-KOH pH 7.6) and 15% v:v LFB1/50 (10% w:v sucrose, 50 mM KCl, 2 mM MgCl_2_, 1 mM EGTA, 2 mM DTT, 20 mM K_2_HPO4/KH_2_PO4 pH 8.0, 40 mM HEPES-KOH, pH 8.0). The extract was clarified by centrifugation at 84,000 x g in a pre-cooled SW55 rotor swinging bucket rotor at 4°C for 20 min. The golden cytoplasmic layer was recovered, supplemented with glycerol to 2% v/v and frozen in aliquots in liquid nitrogen and stored at −80°C until required.

Sperm was recovered from testes isolated from male frogs post mortem following a lethal dose of anaesthetic (0.2% w:v Tricaine mesylate MS222, ∼0.5% w:v NaHCO3, to pH 7.5). Isolated testes were washed carefully to avoid bursting in EB (50 mM KCl, 5 mM MgCl_2_, 2 mM dithiothreitol or β-mercaptoethanol, 50 mM HEPES-KOH, pH 7.6), prior to being finely chopped with a clean razor blade in fresh EB. Recovered lysate was filtered through a 25 µm nylon membrane to remove particulate matter. Filtered sperm was centrifuged at 2,000 x g at 4°C for 5 min; selective resuspension of the sperm pellet allowed separation of the sperm from contaminating erythrocytes; the resuspended sperm was respun and the pellet resuspended in 0.5 ml SuNaSp (0.25 M sucrose, 75 mM NaCl, 0.5 mM spermidine, 0.15 mM spermine, 15 mM HEPES-KOH, pH 7.6) per testis. The sperm was demembranated with the addition of 25 µl per testis lysolecithin (5 mg/ml, in H_2_O) for 10 min at room temperature. Demembranated sperm were respun and resuspended in SuNaSp plus 3% w/v BSA to quench the demembranation reaction. Quenched sperm were respun and resuspended in 100 µl EB plus 30% glycerol per testis, counted using a haemocytometer and stored at −80°C.

Extracts were supplemented with 250 μg/ml cycloheximide, 25 mM phosphocreatine and 15 μg/ml creatine phosphokinase and incubated with 0.3 mM CaCl_2_ for 15 minutes to trigger release from metaphase arrest. For DNA synthesis reactions, demembranated *Xenopus* sperm nuclei were incubated at 3-10 ng DNA/μl in extract. DNA synthesis was assayed by measuring incorporation of [α-^32^P]dATP into acid-insoluble material followed by scintillation counting, as described(Gillespie *et al*., 2012; Gillespie *et al*., 2016). Extract was supplemented with 50 nCi/µl [α-^32^P]dATP from a high activity 10 mCi/ml stock. At the appropriate times 10 µl aliquots were stopped by the addition of 160 µl Stop-C (0.5% w:v SDS, 5 mM EGTA, 20 mM Tris HCl, pH 7.5) plus freshly added 0.2 mg ml Proteinase K (from stock of 20 mg/ml proteinase K, 50% v:v glycerol, 10 mM Tris HCl, pH 7.5) and were incubated at 37°C for 30 min. Samples are precipitated at 4°C for 30 min by the addition of 4 ml 10% TCA (10% w:v TCA, 2% w:v Na_4_P_2_O7.10H_2_O). 40 µl (1% of 4ml) of the total reaction was spotted on a paper disc. Insoluble material was recovered from solution by filtration through a glass fibre filter mounted on a vacuum manifold. The glass fibre filters were twice washed in 5% TCA (5% w:v TCA, 0.5% w:v Na_4_P_2_O7.10H_2_O), once in 100% ethanol and then air dried. The paper and glass fibre filters were then quantified by scintillation counting. Precipitated material was expressed as a percentage of total counts (%TC) from which DNA replication (ng/µl) was calculated by multiplying by a factor of 0.654. The extent of nuclear formation was followed under the light microscope (phase contrast). All incubations were carried out at 23°C.

Licensing factor extract (LFE) was prepared as described(Chong et al., 1997). The initial steps for preparing metaphase extracts were followed. Before the final spin the extract was activated with 0.3 mM CaCl_2_ for 15 minutes then diluted 5 fold with ‘Licensing Factor Buffer’ (LFB: 40 mM Hepes KOH pH 8.0, 20 mM K_2_HPO_4_/ KH_2_PO_4_ pH 8.0, 2 mM MgCl_2_, 1 mM EGTA, 2 mM DTT, 10% (w/v) sucrose and 1 µg/ml each of leupeptin, pepstatin and aprotinin) supplemented with 50 mM KCl (i.e. LFB1/50) and spun to remove membrane components at 84,000 x g in a pre-cooled SW55 rotor swinging bucket rotor at 4°C for 40 mins. The clarified supernatant was frozen, in aliquots, in liquid nitrogen and stored at −80°C till required.

Membrane free extract (MFE) used for TopBP1 immunoprecipitation experiments was prepared as per LFE, above, but was spun at 186,000 x g in a precooled TLA-100 fixed angle rotor at 4°C for 20 mins and recovered supernatant was used immediately.

Immunodepleted extract was prepared as described (Chong *et al*., 1997). Briefly, rProtein A agarose beads (GEHC), preincubated with 2 volumes of serum, were twice incubated with interphase whole egg extract at a ratio of 1 volume extract plus 0.7 volume beads. Twice depleted extract, recovered from the beads, was frozen in aliquots in liquid nitrogen and stored at −80°C.

Immunoprecipitation was carried out as described using affinity purified antibody saturated Dynabeads following the manufacturer’s instructions (Thermo Fisher Scientific).

#### Chromatin Isolation

All chromatin isolations from *Xenopus* egg extracts were undertaken in low adhesive Eppendorf tubes, as described (Gillespie *et al*., 2012; Gillespie *et al*., 2016). Briefly, reactions were stopped by the addition of 400 μl of ice-cold NIBTX (50 mM KCl, 50 mM Hepes KOH pH 7.6, 5 mM MgCl_2_, 2 mM DDT, 0.5 mM spermidine 3HCl, 0.15 mM spermine 4HCl, 0.1% Triton X-100). This was underlayered with 100 μl 15% sucrose in NIBTX. The tubes were spun at 6000 x g for 5 min at 4°C in a swinging bucket rotor. The buffer above the sucrose cushion was removed and the surface of the cushion washed with 200 μl NIBTX before removing the cushion down to ∼15 μl. The tubes were then spun at 13000 x g for 2 min in a fixed angle rotor to focus the chromatin pellet and following this, all the buffer was removed. The chromatin pellet was then resuspended in loading buffer and was subjected to immunoblotting by standard techniques using 4–12% Bis-Tris gradient SDS–PAGE gel (Invitrogen) with the exception of figure 7a for which a 3-8% Tris-Acetate gel (Invitrogen).

For nuclear transfer experiments, 90 μl interphase extract supplemented with appropriate inhibitors was incubated with 26 ng/μl of sperm nuclei for 1 h. Extract was diluted with 1 ml of NIB50 buffer (50 mM KCl, 50 mM Hepes KOH pH 7.6, 5 mM MgCl_2_, 2 mM DDT, 0.5 mM spermidine 3HCl, 0.15 mM spermine 4HCl) without Triton X-100, underlaid with 100 μl NIB50 containing 15% sucrose, and 5 μl NIB50 containing 30% glycerol, and spun at 2100 x g, 5 minutes at 4 °C in a swinging bucket rotor bench centrifuge. The sucrose cushion was washed with 200 μl NIB50, the buffer was removed leaving ∼15 μl. Nuclei were gently re-suspended and frozen in liquid N_2_.

For the geminin chromatin transfer assay (Ferenbach et al., 2005), 80 μl of extract were activated and incubated for 5 minutes with 120 nM recombinant geminin^DEL^, 20 ng/μl of demembranated sperm nuclei were added to the extract and incubated for 12 minutes. Chromatin was isolated by diluting the reaction with 1 ml of NIB50, underlaid with 100 μl of NIB50 containing 15% sucrose, underlaid with 5 μl of NIB50 containing 30% glycerol and spun at 6000 x g, 5 minutes at 4°C in a swinging bucket rotor bench centrifuge. The buffer above the sucrose cushion was washed with 200 μl NIB50 without Triton and the buffer was removed leaving ∼15 μl sample. Chromatin was gently re-suspended by inversion and frozen in aliquots in liquid N_2_.

#### Immunoblotting

Samples were prepared in 6x Laemmli sample buffer heated at 99°C for 4 minutes, and separated on a 4-12% BisTris NuPage gels (Invitrogen) in 1x MOPS running buffer (Invitrogen) or 3-8% Tris-Acetate gels (Invitrogen) in Tris-Acetate running buffer (Invitrogen). Proteins were transferred onto nitrocellulose membrane (GE Healthcare) and images captured using an Odyssey SLX LiCor Scanner (Licor Bioscience) when using fluorescently labelled secondary antibodies or transferred onto PVDF (GE Healthcare) and detected using enhanced chemiluminescence detection (SuperSignal West Pico Chemiluminescent, Thermo Scientific 34087). Images were captured using either an ImageQuant LAS4000 (GEHC) CCD.

For protein visualisation gels were stained with either Sypro Ruby and de-stained with 10% ethanol, 7% acetic acid, in H_2_O or with Coomassie R-250 and de-stained with 40% ethanol, 10% acetic acid in H_2_O. Quantification of the immunoblots were carried out using pre-diluted protein assay standards (Thermo Scientific 23213), and analysed using Image Studio Lite LICOR software.

#### Mass Spectrometry

Immunoprecipitated samples for mass spectrometry were run on NuPage gels and stained with SimplyBlue SafeStain (Invitrogen). Bands at the size of Treslin and MTBP were cut from gels and reduced with dithiothreitol, alkylated with iodoacetamide and in-gel digested with trypsin. The extracted peptide solutions were analysed using nano LC-MS/MS on an LTQ Orbitrap Velos (ThermoFisher, San Jose, CA).

#### Gel Filtration and Glycerol Gradients

For gel filtration, 100 µl of a 3% PEG precipitate re-suspended in LFB1/50 to a final volume of twice that of undiluted extract was applied to an Agilent BIO SEC-5 (500 Å 4.6 x 300 mm Range 15-5000 kDa), equilibrated in LFB1/50. The column was run at a flow rate of 150 μl/min on an UltiMate3000 chromatography system (Thermo Scientific), at 4°C and 75 μl fractions were collected. For sucrose gradient centrifugation, 440 µl of a 3% PEG precipitate re-suspended in LFB1/50 to a final volume of twice that of undiluted extract was applied to an 11 ml 5-40% sucrose gradient prepared in LFB1/50. Gradients were spun at 40,000 rpm (∼111,000 g) for 20 h in a swinging bucket SW41Ti rotor, in a Beckman Optima ultracentrifuge. 440 μl fractions were collected. Molecular masses were calculated according to Siegel and Monty (Siegel and Monty, 1966) using the values: thyroglobulin tetramer (1338 kDa, 107 Å), thyroglobulin dimer (669 kDa, 85 Å, 19.5 S), apoferritin (443 kDa, 67 Å, 17.6 S), β--amylase (200 kDa, 54 Å, 8.9 S), BSA (66 kDa, 35.5 Å, 4.3 S), carbonic anhydrase (29 kDa, 24.3 Å, 3.2 S) (Sigma-Aldrich MWGF1000).

#### Partial purification of Treslin-MTBP

PEG precipitation was carried out as described (Chong *et al*., 1997). Briefly, to one volume of extract, the required amount of 50% v: v of polyethylene glycol (PEG) 6000 in H_2_O was added, mixed well, and incubated for 30 mins on ice-water. The samples were spun at 20000 g, 30 minutes at 4 °C in a fixed angle rotor. The supernatant was removed and the pellet re-suspended in the required amount of LFB1/50 (50 mM KCl, 20 mM K_2_HPO_4_/KH_2_PO_4_ pH 8.0, 40 mM HEPES–KOH, 2 mM MgCl_2_, 1 mM EGTA, 2 mM dithiothreitol, pH 8.0).

SP and ANX Hi-trap 1 ml columns (Cytiva LifeSciences) were washed with 5 column volumes of H_2_O, 5 column volumes of LFB1/50, 5 column volumes of LFB1/50 containing 1 M KCl and 10 column volumes of LFB1/50. The samples were spun at 21910 g for 20 mins at 4°C in a fixed angle rotor on a bench centrifuge before loading. Columns were connected to an AKTApurifier UPC10 system (Cytiva LifeSciences) or to an UltiMate3000 chromatography system (Thermo Scientific). When the OD_280_ was flat the proteins bound to the matrix of the columns were eluted through a gradient from 50 to 750 mM KCl in 14 x 1 ml fractions at a flow rate of 1 ml/minute.

#### Quantification And Statistical Analysis

Statistical data for individual experiments are presented in the appropriate figure legends. In all cases ‘n’ is the number of independent experimental repeats from which the mean ±S.E.M. has been calculated.

