## Supplemental figure 1 for "The role of DDK and Treslin-MTBP in coordinating replication licensing and pre-Initiation Complex formation"

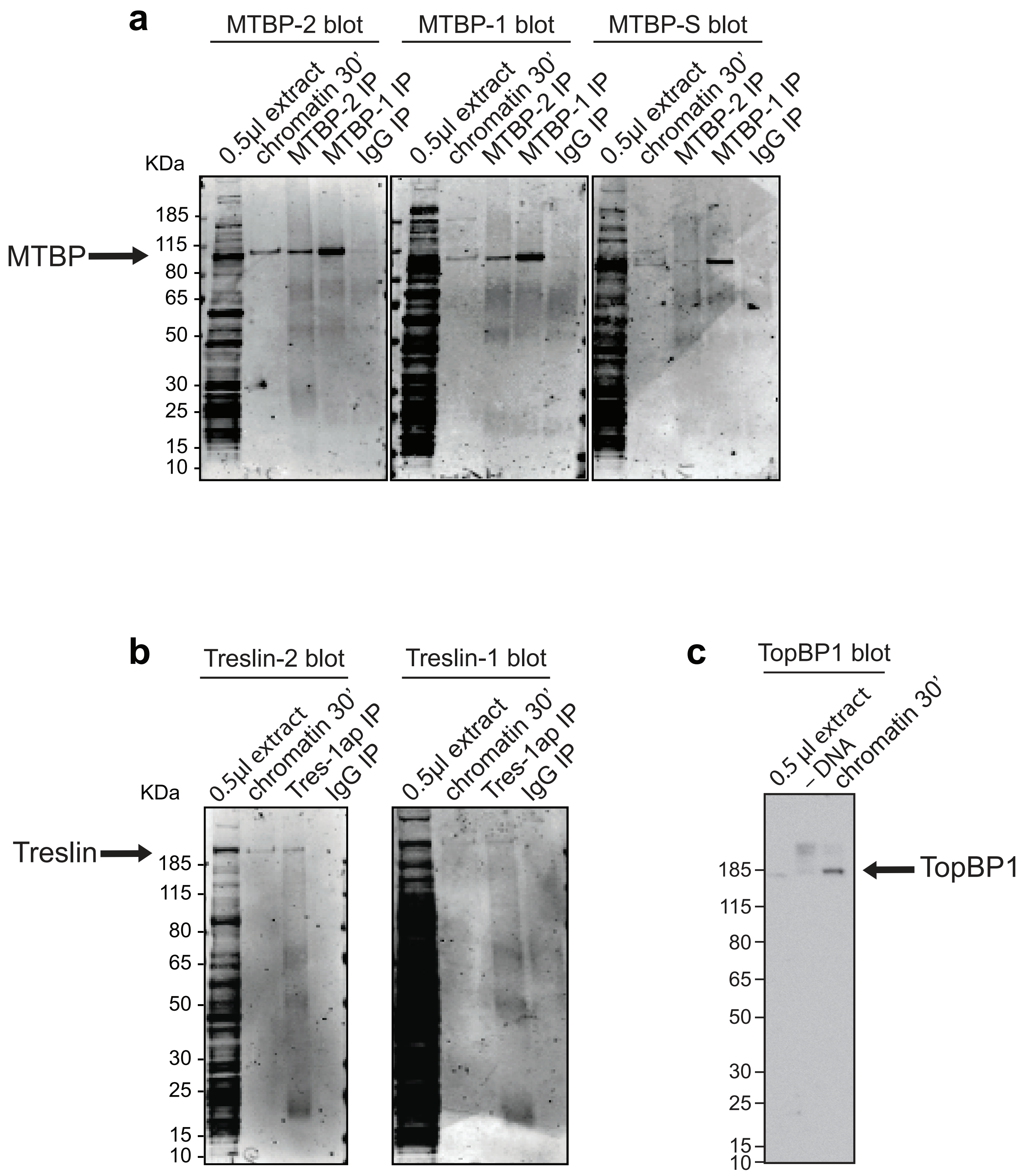


**Figure S1. Test of MTBP, Treslin and TopBP1 antibodies.**
**a.** Interphase extract, S-phase chromatin and IPs carried out using MTBP-1 affinity-purified (MTBP-1 IP), MTBP-2 affinity-purified (MTBP-2 IP) and IgG (IgG IP) antibodies were immunoblotted using affinity-purified MTBP-1, MTBP-2 and MTBP-S (anti human MTBP from Sigma-Aldrich). **b.** Interphase extract, S-phase chromatin and IPs carried out using Treslin-1 affinity-purified (Tres-1ap IP) and IgG (IgG IP) antibodies were immunoblotted using protein-A purified Tres-2ap and affinity-purified Treslin-1 antibodies. **c.** Interphase extract, S-phase chromatin and mock chromatin preparation from extract lacking DNA (-DNA) blotted with affinity-purified TopBP1 antibody. **c.** Whole extract, or chromatin prepared from extract incubated for 30 mins plus or minus sperm DNA was blotted with TopBP1 antibodies.
