## Supplemental figure 2 for "The role of DDK and Treslin-MTBP in coordinating replication licensing and pre-Initiation Complex formation"

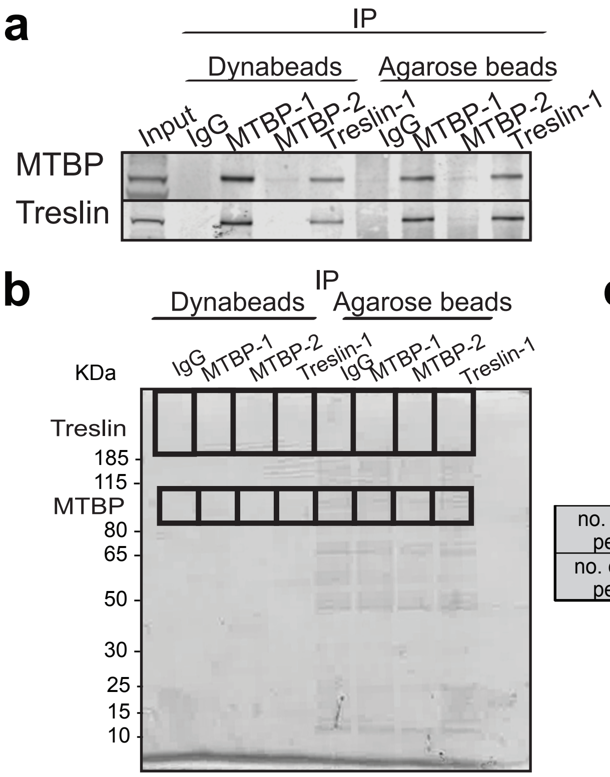


c

| | IP Agarose beads | | | | IP Dynabeads | | | |
|---|---|---|---|---|---|---|---|---|
| | IgG | MTBP1 | MTBP2 | Treslin | IgG | MTBP1 | MTBP2 | Treslin |
| no. of MTBP peptides | 4 | 41 | 8 | 33 | 0 | 47 | 10 | 33 |
| no. of Treslin peptides | 0 | 16 | 1 | 23 | 0 | 16 | 0 | 4 |

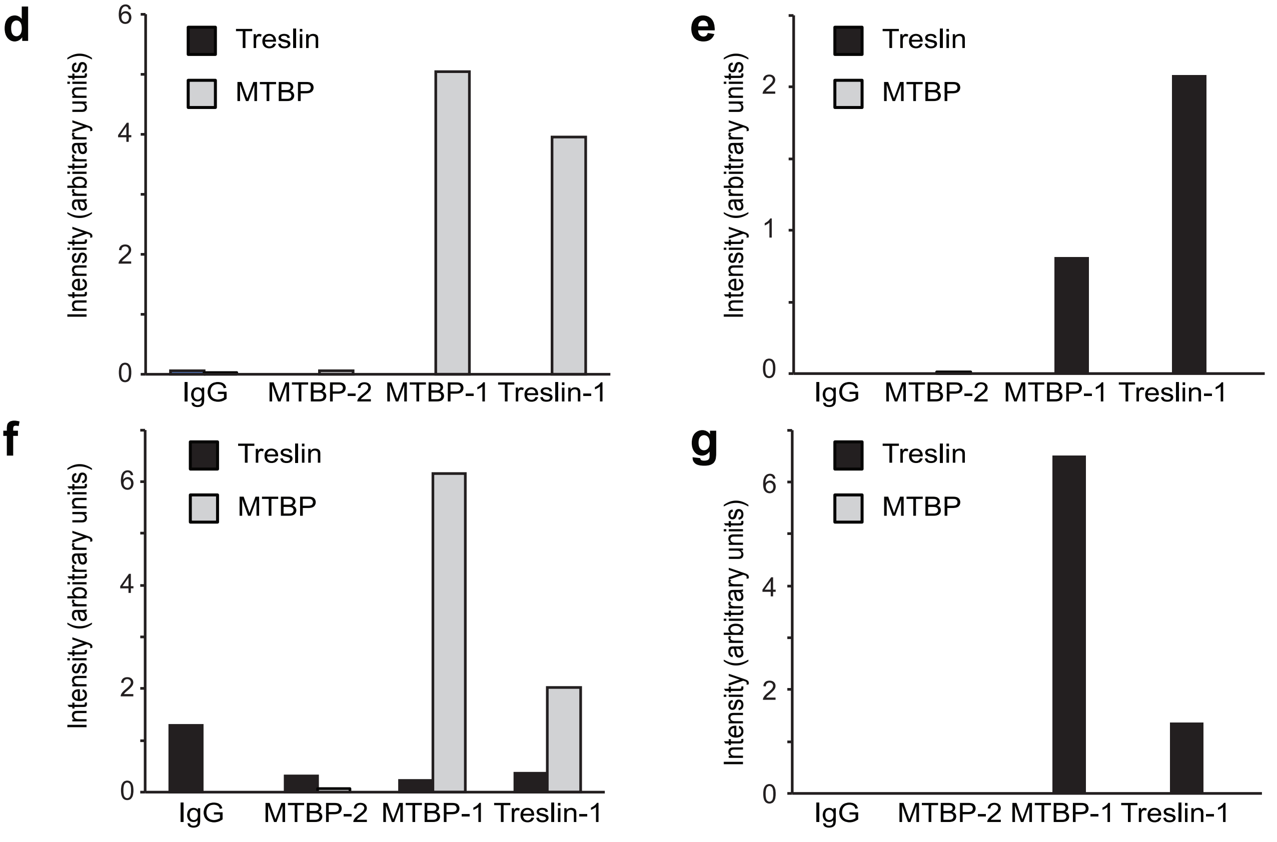


**Figure S2. Mass spectrometry of immunoprecipitation carried out using MTBP-1, MTBP-2 and Treslin antibodies.**

Immunoprecipitation using MTBP-1, MTBP-2 and Treslin-1 affinity purified antibodies were carried out from interphase extract using protein-A Dynabeads and agarose beads. **a.** Immunoprecipitates were immuno- blotted for MTBP and Treslin. **b.** Immunoprecipitates were run on NuPage gel and stained with Coomassie. The indicated bands were cut for MS analysis. **c.** Number of MTBP and Treslin peptides counted in IgG, MTBP-1, MTBP-2 and Treslin-1 immunoprecipitates. **d.** Intensity of MTBP and Treslin peptides counted in the IPs carried out with agarose beads, detected in the MTBP-molecular weight bands. **e.** Intensity of MTBP and Treslin peptides counted in the IPs carried out with agarose beads, detected in the Treslin-molecular weight bands. **f.** Intensity of MTBP and Treslin peptides counted in the IPs carried out with Dynabeads beads, detected in the MTBP-molecular weight bands. **g.** Intensity of MTBP and Treslin peptides counted in the IPs carried out with Dynabeads beads, detected in the Treslin-molecular weight bands.
