## Supplemental figure 3 for "The role of DDK and Treslin-MTBP in coordinating replication licensing and pre-Initiation Complex formation"

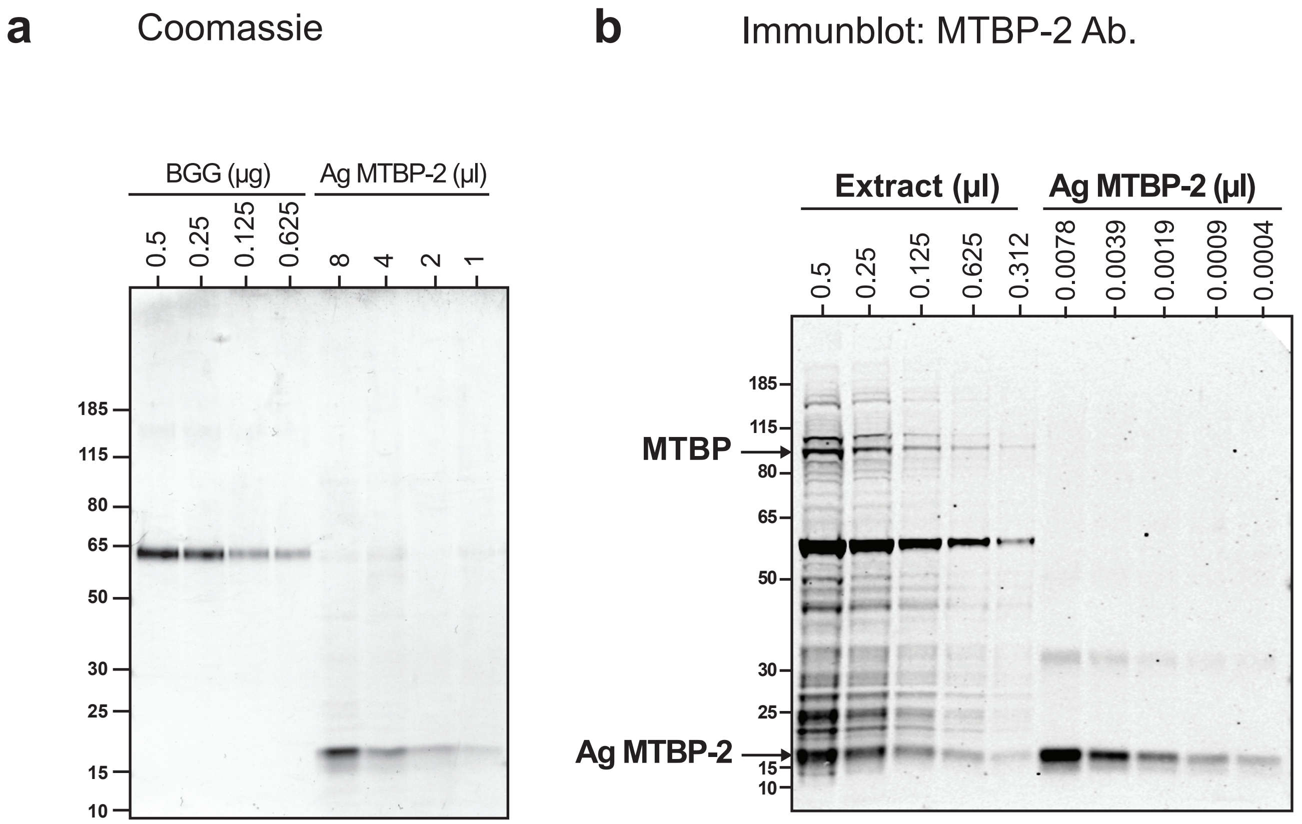


**Figure S3. Quantitation of amount of MTBP in xenopus egg extract.**
**a.** Different amounts of MTBP-2 antigen was loaded on a NuPage gel with a titration of Bovine Gamma Globulin. The gel was stained with Coomassie, and analysed with Image Studio Lite LiCor software for quantification. The average concentration of MTBP-2 antigen derived based on three independent experiments is 27.6 ng/µl or 1.93 µM.
**b.** Different amounts of MTBP-2 antigen along with *Xenopus* egg extract were run on NuPage gel and immunoblotted with MTBP-2 affinity-purified antibody. The average concentration of MTBP in extract derived from known antigen concentration based on three independent experiments is 0.193 ng/µl or 2 nM.
