## Supplemental figure 4 for "The role of DDK and Treslin-MTBP in coordinating replication licensing and pre-Initiation Complex formation"

| Protein names | Gene names | Intensity IgG | Intensity MN |
|---|---|---|---|
| Zinc finger protein gli3 | gli3 | 0 | 219480000 |
| DNA topoisomerase 2A | LOC398512;top2a | 0 | 82799000 |
| Kinesin-like protein KIF2C | kif2c | 0 | 80648000 |
| Mdm2-binding protein | mtbp | 0 | 34621000 |
| Eukaryotic translation initiation factor 2 subunit 1 | eif2s1 | 0 | 17309000 |
| ATP binding cassette subfamily F member 1 | abcf1 | 0 | 12640000 |
| Kinesin-like protein; Chromosome associated kinesin KIF4 | kif4a-A-prov;kif4 | 0 | 10849000 |
| Werner syndrome ATP-dependent helicase homolog | FFA-1;wrn | 0 | 10843000 |
| Chloride channel CLIC-like protein 1 | LOC397947;clcc1 | 0 | 10506000 |
| Receptor protein serine/threonine kinase | XBMPR-II | 0 | 10336000 |
| Orc1 | XORC1;orc1 | 0 | 8280800 |
| Treslin | ticrr | 0 | 7942600 |
| Eukaryotic translation initiation factor 2 subunit beta | eif2s2 | 0 | 6634400 |
| Orc3 | orc3l;orc3 | 0 | 6153700 |
| UPF2, regulator of nonsense mediated RNA decay | upf2 | 0 | 5360200 |
| Integrator complex subunit 2 | ints2 | 0 | 4243800 |
| Inner centromere protein A | incenp-a | 0 | 2526000 |

**Figure Supplementary Figure S4. MTBP-immunoprecipitation from partially purified material.**
List of protein detected by MS specifically in the MTBP-1 immunoprecipitate with intensity values.
