## Supplemental figure 5 for "The role of DDK and Treslin-MTBP in coordinating replication licensing and pre-Initiation Complex formation"

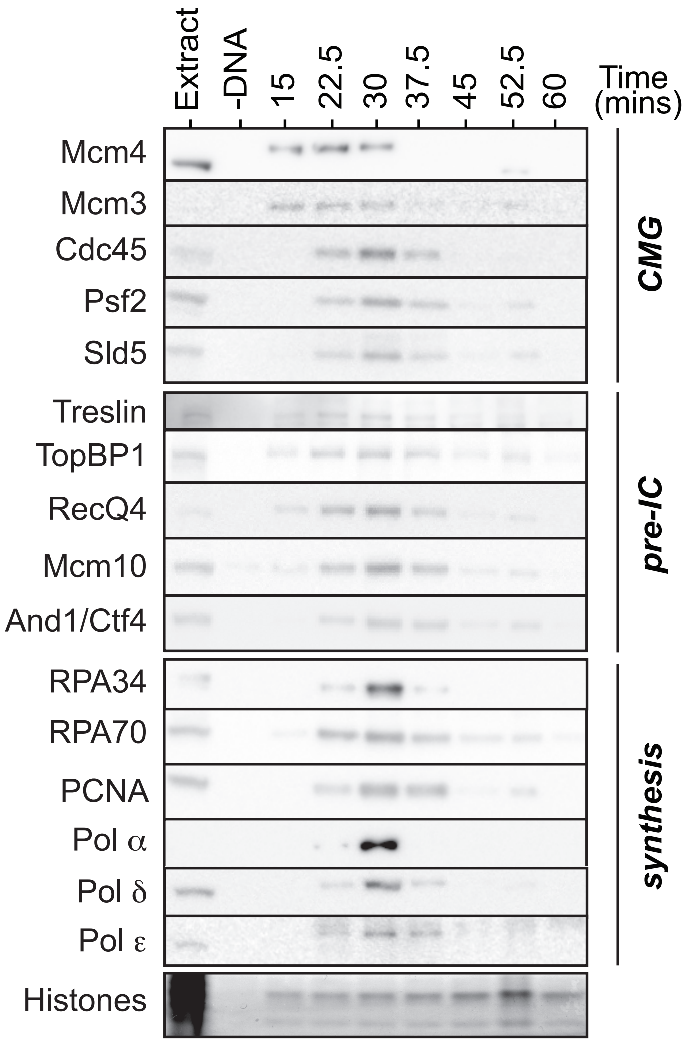


**Figure S5. Replication protein chromatin association timecourse**
Sperm nuclei were incubated in interphase extract for the indicated times; chromatin was isolated, run on a NuPage SDS-PAGE gel and immunoblotted for the indicated proteins. The lower portion of the gel was stained with Coomassie to detect histones.
