## Supplemental figure 6 for "The role of DDK and Treslin-MTBP in coordinating replication licensing and pre-Initiation Complex formation"

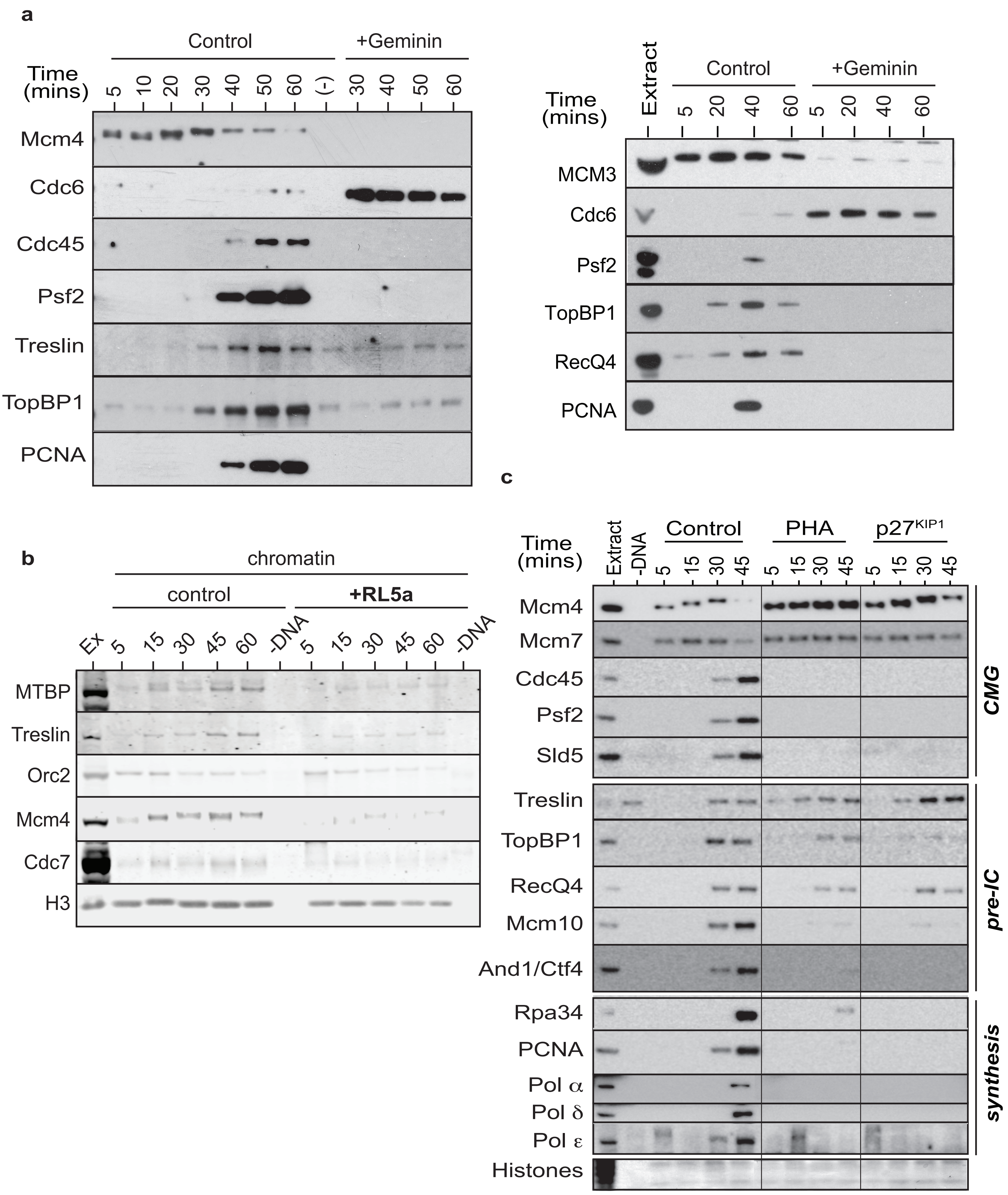


**Figure S6. Replication protein chromatin association timecourse**

Sperm nuclei were incubated in interphase extract for the indicated times ± **a.** geminin, **b.** RL5a and **c.** PHA-767491 or p27KIP1; chromatin was isolated, run on a NuPage SDS-PAGE gel and immunoblotted for the indicated proteins. The lower portion of the gel was stained with Coomassieto detect histones.
