## Supplemental figure 7 for "The role of DDK and Treslin-MTBP in coordinating replication licensing and pre-Initiation Complex formation"

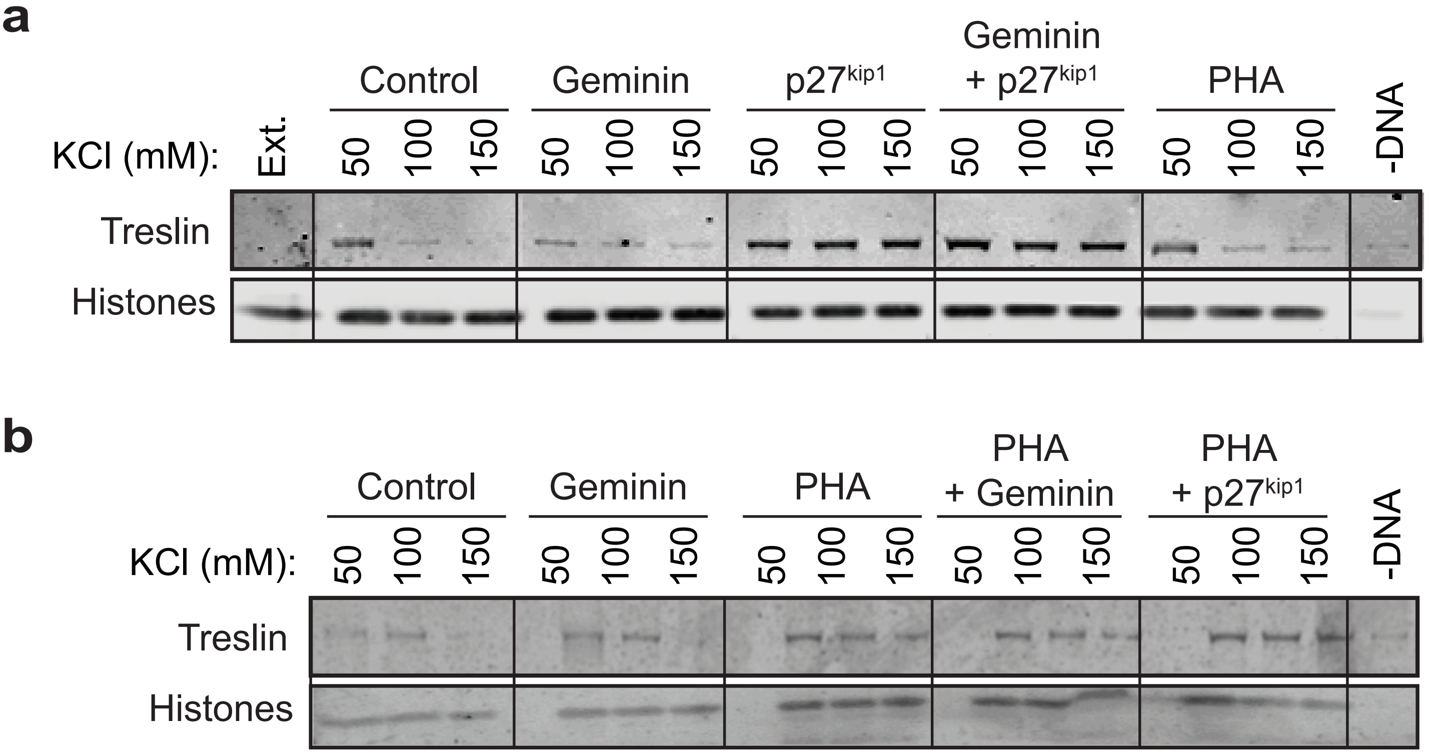


**Figure S7. Salt resistant Treslin chromatin association**

Sperm nuclei were incubated in interphase extract for 60 mins in the presence of either geminin, p27KIP1, PHA or a combination of 2 of these; chromatin was isolated at the indicated salt concentration, run on a NuPage SDS-PAGE gel and immunoblotted for Treslin. The lower portion of the gel was stained with Coomassie to detect histones.
