## Supplemental figure 8 for "The role of DDK and Treslin-MTBP in coordinating replication licensing and pre-Initiation Complex formation"

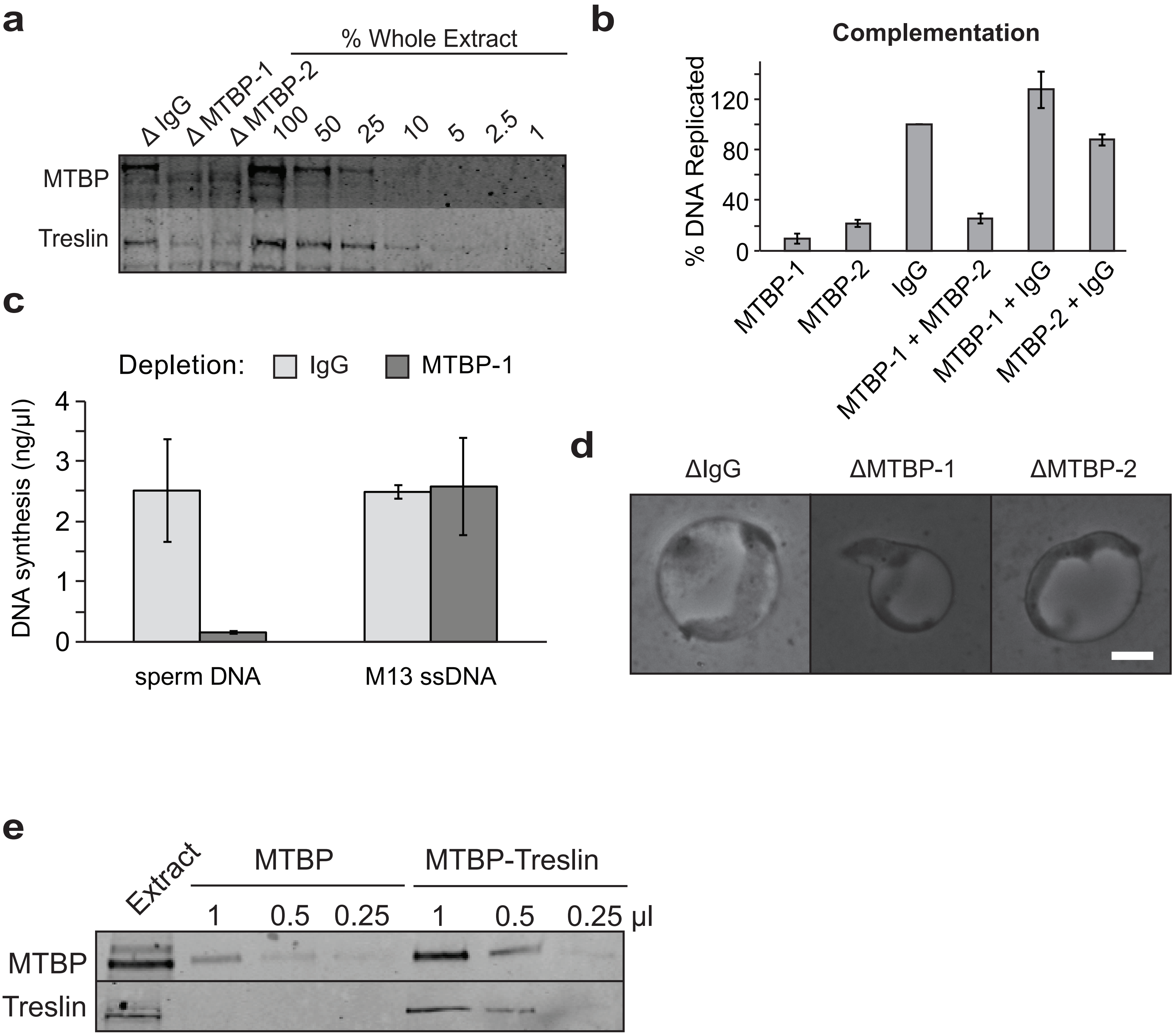


**Figure S8. MTBP immunodepletion. a.** Mock, MTBP-1 and MTBP-2 depleted extracts were run on a NuPage SDS-PAGE gel together with a titration of whole egg extract and immunoblotted for MTBP and Treslin. **b.** An equal amount of mock, MTBP-1 and MTBP-2 depleted extract were mixed, sperm nuclei were added and the extent of DNA replication was determined after three hours. (100% = replication in mock depleted extract). Average of three independent experiments, error bars represent standard error. **c.** Either sperm nuclei or M13 ssDNA were incubated for three hours in mock or MTBP-depleted extracts and the extent of DNA replication was determined. Average of three independent experiments, error bars represent standard error. Sperm nuclei and M13 ssDNA were added to a final DNA concentration of 3 ng/μl. **d.** Phase contrast images of nuclei formed after one hour of incubation in the indicated depleted extract. Scale bar represents 25 μm. **e.** Titration of fractions containing MTBP or MTBP-Treslin were run on a NuPage SDS-PAGE gel and immunoblotted for MTBP and Treslin.
