## Supplemental figure 9 for "The role of DDK and Treslin-MTBP in coordinating replication licensing and pre-Initiation Complex formation"

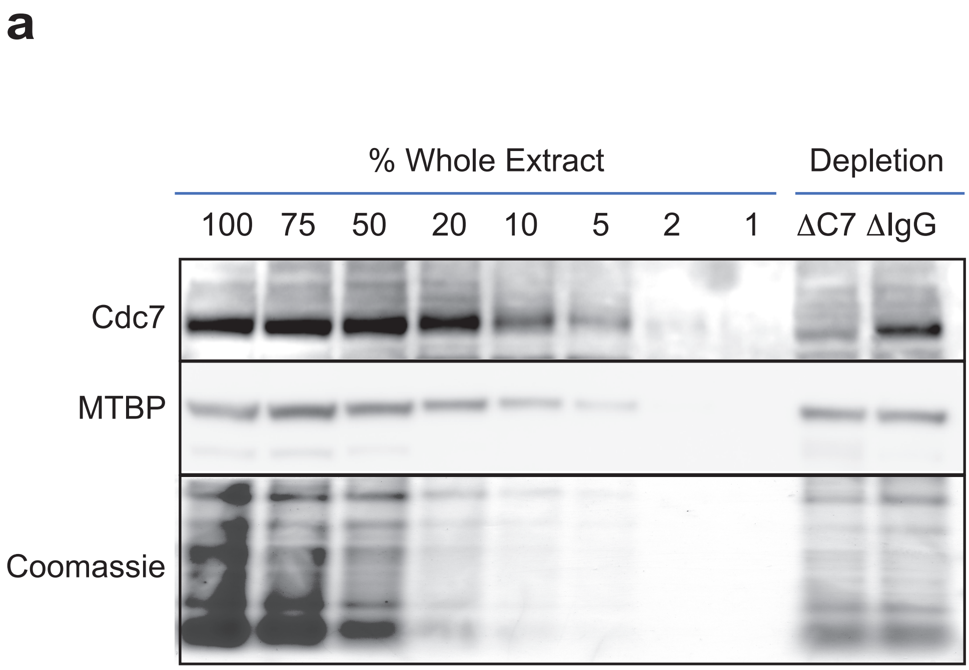


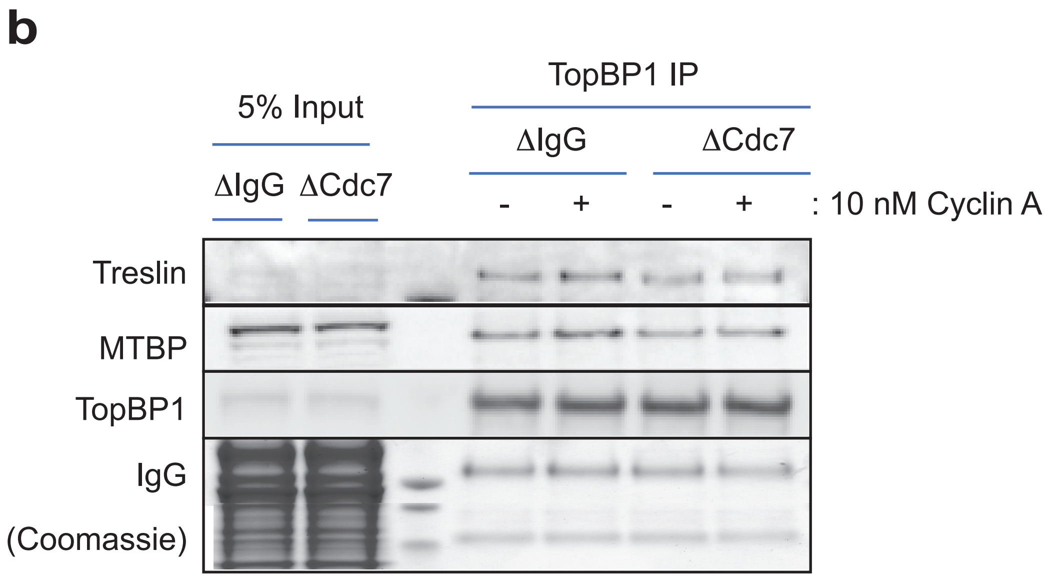


**Figure S9. Treslin-MTBP-TopBP1 complex formation**

**a.** interphase extract was immunodepleted using either control serum or anti-Cdc7 serum. Recovered extract was run on a NuPage SDS-PAGE gel and immunoblotted for the indicated proteins, together with a titration series of whole egg extract to determine the extent of extract depletion. **b.** Extract depleted with either controll serum or anti-Cdc7 (DDK) antobody was either spplemented or not with 10 nM cyclin A, as indicated and incubated for 15 mins. Samples were immunoprecipitated using either control IgG or anti-TopBP1-antibody bound protein-G DynaBeads. Washed precipitates were subjected to SDS-PAGE on a NuPage gel. The upper portion of the gel was immunoblotted for Treslin, MTBP and TopBP1 as indicated and the lower portion of the gel was stained with Coomassie to visualise the Heavy and Light chains of IgG. Extract samples, 5% of input for the IP are shown.
